# Flexible regulation of representations on a drifting manifold enables long-term stable complex neuroprosthetic control

**DOI:** 10.1101/2023.08.11.551770

**Authors:** Nikhilesh Natraj, Sarah Seko, Reza Abiri, Hongyi Yan, Yasmin Graham, Adelyn Tu-Chan, Edward F. Chang, Karunesh Ganguly

## Abstract

The nervous system needs to balance the stability of neural representations with plasticity. It is unclear what is the representational stability of simple actions, particularly those that are well-rehearsed in humans, and how it changes in new contexts. Using an electrocorticography brain-computer interface (BCI), we found that the mesoscale manifold and relative representational distances for a repertoire of simple imagined movements were remarkably stable. Interestingly, however, the manifold’s absolute location demonstrated day-to-day drift. Strikingly, representational statistics, especially variance, could be flexibly regulated to increase discernability during BCI control without somatotopic changes. Discernability strengthened with practice and was specific to the BCI, demonstrating remarkable contextual specificity. Accounting for drift, and leveraging the flexibility of representations, allowed neuroprosthetic control of a robotic arm and hand for over 7 months without recalibration. Our study offers insight into how electrocorticography can both track representational statistics across long periods and allow long-term complex neuroprosthetic control.

## 1. Introduction

Our nervous system needs to balance maintaining a stable neural representation of a large repertoire of well-rehearsed actions while also facilitating new learning. We use the term ‘representation’ here to refer to network-wide cortical activity patterns during repeated performance of a large repertoire of actions. Studies in animals have indicated that the neural representations of behavior can experience drift – large-scale changes in the correlation between neural activity and behavior over days and weeks (Clopath et al., 2017; Rule et al., 2019; Schoonover et al., 2021). While past work in humans using functional imaging as well as invasive recordings have indicated that simple well-rehearsed movements that are part of daily life, such as finger flexion and tongue protrusion, have distinct somatotopic representations on a given day (Lotze et al., 2000; Leuthardt et al., 2004; Miller et al., 2007; Meier et al., 2008; Schellekens et al., 2018; Degenhart et al., 2018), it remains unclear how such representations change over days and weeks. Especially given the significantly longer lifespan of human beings and the extent of repetitions, it is not clear whether a repertoire of simple actions may not demonstrate drift. Moreover, how new contextual motor learning might affect representational dynamics is unclear.

A key challenge has been to accurately track over time, and with high spatiotemporal resolution, the neural representations during the performance of a repertoire of simple actions. We used intracortical brain-computer interfaces (BCIs) based on mesoscale electrocorticography (ECoG) (Leuthardt et al., 2004; Wang et al., 2013; Moses et al., 2021; Silversmith et al., 2021; Benabid et al., 2019; Degenhart et al., 2018; Angrick et al., 2023, 2021; Vansteensel et al., 2016; Pels et al., 2019; Metzger et al., 2022) to understand principles of how the nervous system might balance the representational stability and plasticity of a repertoire of simple, imagined actions spanning the whole body. We first established that an ECoG grid covering large-scale sensorimotor cortex across a single brain hemisphere could represent imagined movements of body parts spanning the whole body. This allowed us to identify a representational structure based on the pairwise separation or distance between movements (Natraj et al., 2022; Ejaz et al., 2015; Leonard et al., 2020) on an across-day preserved low-dimensional mesoscale ‘manifold’ (Natraj et al., 2022; Ottenhoff et al., 2022; Flint et al., 2020; Bouchard et al., 2013; Gallego et al., 2017). We then assessed the neural statistics affecting the representational structure across days and during long-term learning in the context of complex neuroprosthetic control.

We found that the mesoscale representations of each imagined action – ‘open-loop’ and without feedback – demonstrated a net drift across days. Specifically, the centroids of their respective distributions drifted; this also resulted in net shifts of the manifold location across days. However, the relative representational structure across the repertoire on the manifold was remarkably stable day-to-day. Strikingly, variance within and across repeated presentations could be flexibly modulated during closed-loop BCI control - with visual feedback - to stably modify the representational structure. Such representational changes were most pronounced after closed-loop decoder adaptation. Thus, within any single BCI session in comparison to the open-loop condition, there were rapid increases in discernability between movements. This was driven primarily by variance reductions and notably without any relative changes in somatotopy. With such feedback and practice, discernability between the representation of actions increased across days and remained remarkably contextually specific to the BCI. From a translational perspective, accounting for drift in our decoders and leveraging the flexibility of representations across BCI contexts allowed ‘plug and play’ control – i.e., without any need for decoder updates (Silversmith et al., 2021). This culminated in skilled reach-to-grasp neuroprosthetic control for over 7 months with a fixed deep-learning based decoder. Our findings indicate that ECoG signals can offer fundamental insight into the relative representational stability of a repertoire of well-rehearsed actions in humans as well as how their statistics can be adapted, despite net representational drift, for novel contextual learning. This also enabled highly clinically relevant long-term stable neuroprosthetic control, resembling the consolidation of skilled movement control in able-bodied subjects.

## 2. Results

### 2.1. ECoG recordings of imagined whole-body movements

We recorded 128-channel electrocorticographic activity (ECoG) from sensorimotor cortex in two subjects (Fig. 1A) as part of the ongoing BRAVO clinical trial at UCSF (NCT03698149). The first subject (B1) is a right-handed male with severe tetraparesis and anarthria due to a brain-stem stroke. The second subject (B2) was a right-handed male with end-stage ALS and completely paralyzed, from whom we had a limited window to record ECoG activity. The grid of ECoG channels was placed on the hemisphere contralateral to the dominant hand. Raw activity was filtered, and power was computed in three canonical sensorimotor frequencies of interest: *δ* - 0.5 to 4Hz, *β* - 12 to 30Hz and *γ_H_* - 70 to 150Hz (Silversmith et al., 2021; Natraj et al., 2022). Subjects were instructed to imagine performing whole-body movements (Willett et al., 2020; Degenhart et al., 2018; Hochberg et al., 2006) in a trial-based design (Fig. 1A-B). Trial averaged M1 (primary motor cortex) *γ_H_* activity during the active period of the task (Fig. 1C and 1D, z-scored relative to a baseline period) revealed significant activity for whole-body movements. In B1 with whom we were able to capture a wider repertoire of movements, the magnitude of z-scored *γ_H_* activity in M1 was suggestive of a somatotopic map (Fig. 1C). Indeed, *γ_H_* event-related potentials (ERPs) highlighted spatially distinct activity patterns for the movement repertoire. This is shown in Fig. 1E for face-centric movements and in Fig. 1F for the hand (see Fig. S1 for examples of *δ* and *β* ERPs). Identifying channels across the grid with significant ERPs (*p* ≤ 5 × 10^−4^) further revealed differing body-centric somatotopic maps in spatial activity patterns (*γ_H_* maps in Fig. 1G).

**Figure 1:**
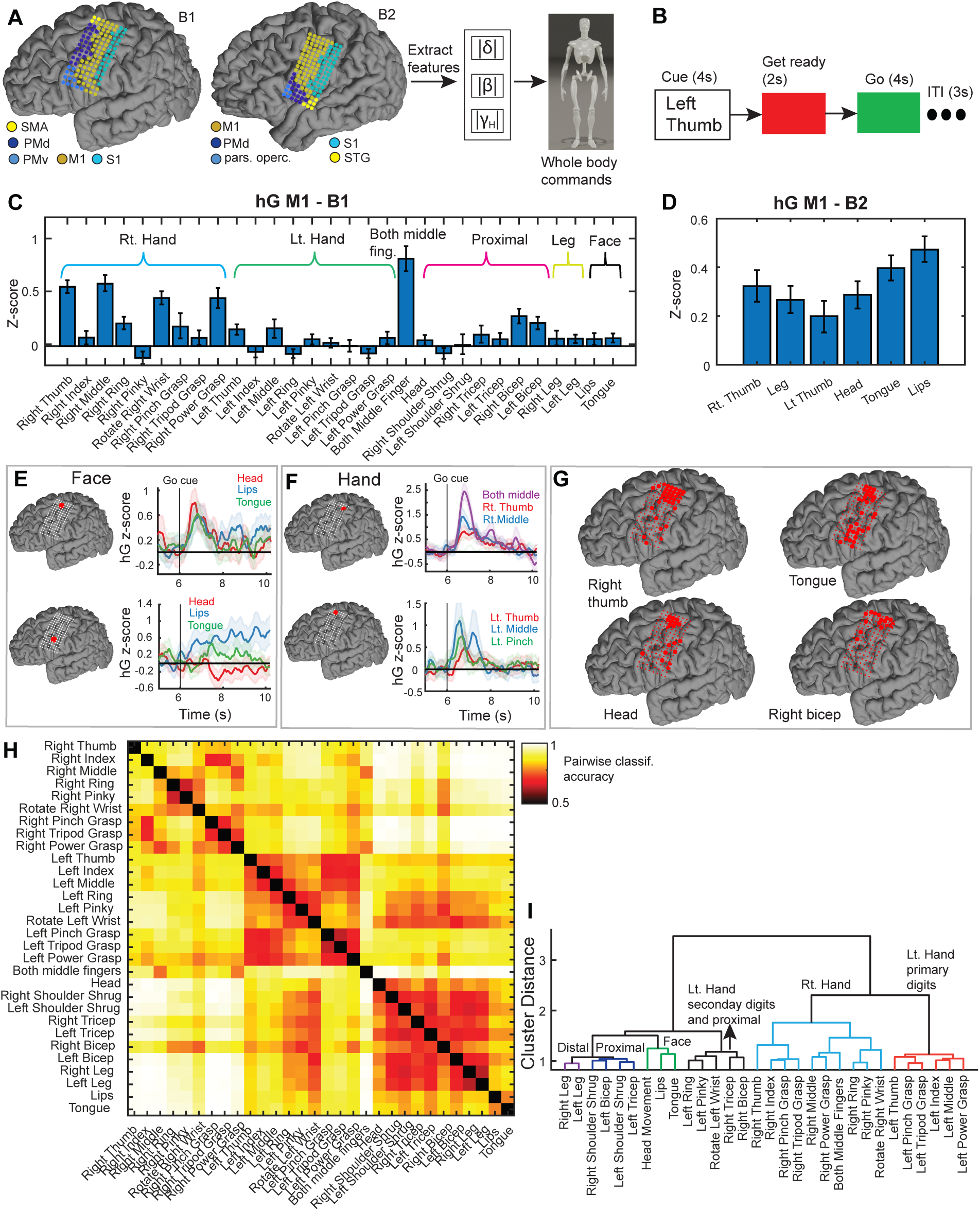
Whole-body somatotopic representation of imagined movements from a unihemispheric ECoG grid over sensorimotor cortex. A) ECoG grid placement with electrodes color-coded by anatomical location for the two subjects. Neural features in three canonical frequencies of interest were extracted as subjects imagined performing body part movements. B) Event-related task design when collecting imagined movements across the whole body (ITI – inter trial interval). C-D) *γ_H_* activity (mean and S.E., z-scored relative to a baseline period) in primary motor cortex channels (M1) in subjects B1 and B2 during imagined movements. E-F) *γ_H_* event related potentials (ERPs) for Face-centric and Hand-centric movements in B1 (mean and shaded S.E.M.) for the trial design in Fig. 1B, at two exemplar electrodes (red channels). The black vertical line indicates the appearance of the Go cue. G) *γ_H_* somatotopy for representative movements in B1, highlighting spatially distinct channels for different movements based on significant ERPs. H) Cross-validated pairwise classification accuracies between movements’ neural distributions across all channels and frequency bands for B1 during the active portion of the task in Fig. 1B. Values closer to 1 imply dissimilarity and closer to 0.5 imply chance levels of separation. I) Hierarchical clustering of the matrix in Fig. 1H revealed body part-specific similarity in spatial activity patterns i.e., somatotopic coding.

We further examined the structure of somatotopic activity patterns in B1 by comparing spatial activity across the grid in all three frequency bands for the repertoire (Methods). Thsi revealed a somatotopic, body-centric representational structure (Fig. 1H and 1I) in B1. Divergent effectors (e.g., hand vs. feet) tended to have distinct neural activity whereas within-effector movements (e.g., digits of the hand) tended to be more like each other. Parcellation of right-handed movements revealed a grasp vs. finger-specific scheme with a unique representational structure similar to overt movements in able-bodied subjects (Natraj et al., 2022). Notably, left-handed movements clustered with the right limb based on the digits being controlled; primary digit movements (left thumb, index, middle) were more similar to right-handed movements whereas secondary digits (pinky, ring) were more like proximal right arm movements. Overall, our analyses provided evidence of a somatotopic representational structure across the entire body on the sensorimotor cortex of a single hemisphere.

### 2.2. Design of the BCI experiments and decoders

For the BCI experiments, we selected a subset of the discrete movements and mapped each movement onto an axis in virtual 3D space (including the origin) for egocentric-based con-trol of a a virtual Jaco robotic arm and hand (Kinova Assistive Robotics) (Fig. 2A, B1). In B2, where we had a more limited opportunity to collect data, we mapped four movements onto the four cardinal directions of a 2D workspace (Fig. S2A). In both subjects, these mappings were done while retaining a body-centric reference scheme, i.e., virtual movements in the Y direction corresponded to respective imagined head (positive) and feet (negative) actions. We measured the representational structure of the sub-selected movements across three experiments: 1) in the open-loop experiment (OL), subjects were instructed to imagine performing the movement associated with a cued cardinal direction (green square, 2B). They did not receive any visual feedback of their mental state, providing a window into open-loop sensorimotor representations. This open-loop data was used to seed a discrete real-time neural classifier (Methods). 2) In the first closed-loop experiment (CL1), the subject now received real-time visual feedback of the classifier output via a red arrow (Fig. 2C, right) pointing in the respective axis (X, Y, Z) associated with the decoded movement class. The visual feedback allows the subject to adapt their sensorimotor representations online to correctly orient the arrow (Fig. 2C). 3) The second closed-loop experiment (CL2) followed the same procedure as CL1, with the difference that the weights of the decoder were updated with CL1 data. To update the weights, we assumed in CL1 that the user always intended to direct the decoded arrow toward the cued target (Methods, (Gilja et al., 2012)). Together, CL1 and CL2 provide a window into how the sensorimotor representational structure adapts to two BCI-specific contexts.

**Figure 2:**
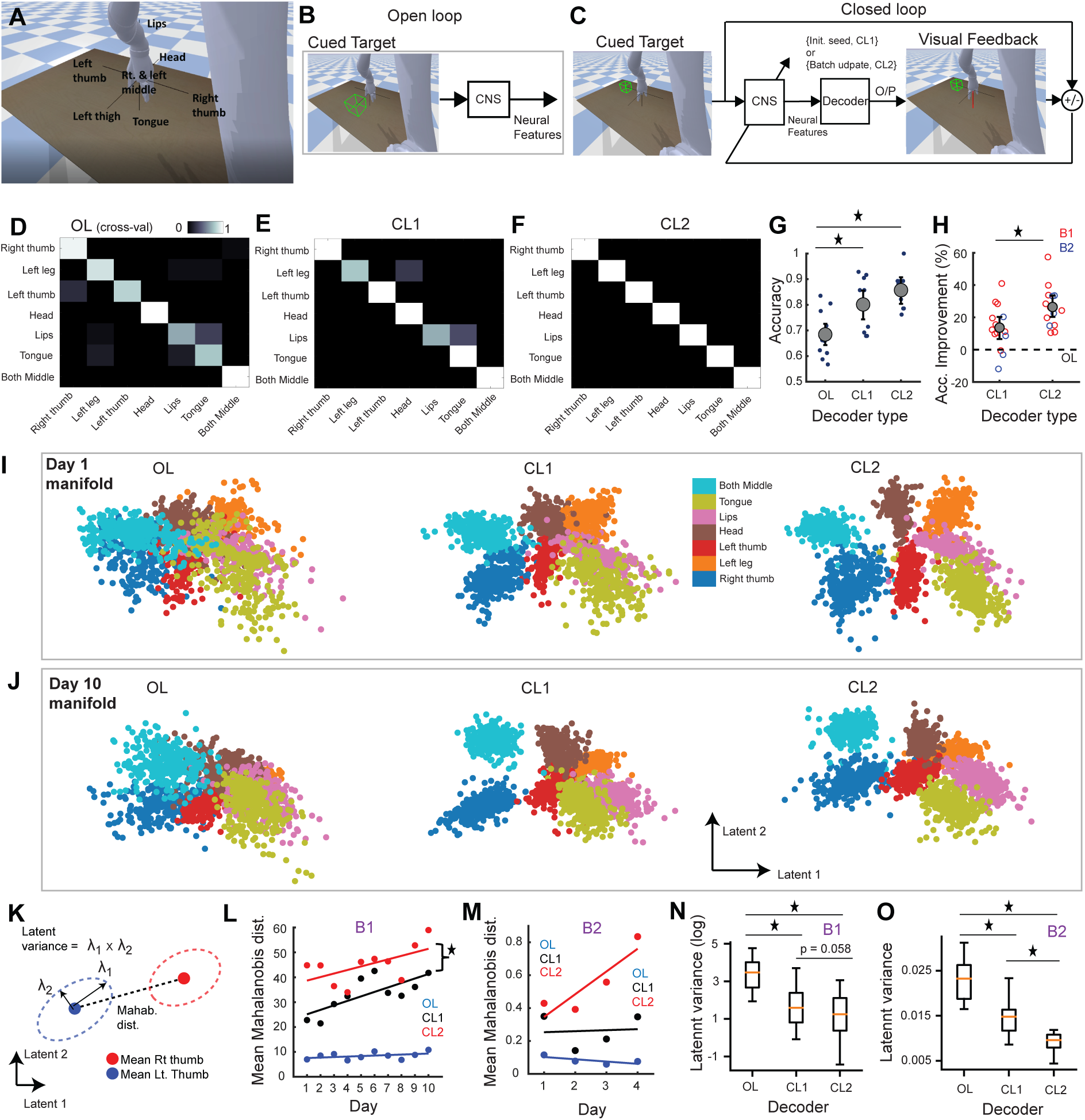
BCI-specific short and long-term plasticity of a stable open-loop representational structure. A) Assigning individual imagined movements as discrete neural commands to control a virtual effector along the cardinal 3D axes. B) Design of the open-loop experiment (OL) where the subject imagined performing the action associated with the cued direction but without visual feedback. C) Design of the closed-loop experiments with instantaneous visual feedback. D) Example cross-validated open-loop confusion matrix on predicting the movement class from held-out trials in B1. E-F) Example CL1 and CL2 confusion matrix from online trials in B1. G) Average decoding accuracy (with S.E.) in OL, CL1, and CL2 experiments in B1. Each dot represents performance in an individual day. The star sign represents a significant difference in means between variables grouped by the horizontal line. (∗) : *p* ≤ 0.05 here and after. H) Improvements in decoding accuracy during CL1 and CL2 each day relative to OL performance (black dotted line). Each dot represents performance at an individual session color-coded by the subject. The star sign represents a significant difference in means between variables grouped by the horizontal line. I-J) Neural samples from Day 1 and Day 10 projected onto their own latent-space for the three different experiments in subject B1, color-coded by the class of movement. K) Representational distance between any two movements (in 2D latent-space) via the Mahalanobis distance, i.e., distance between means scaled by the inverse of the pooled covariance matrix. The neural variance of each movement was computed as the product of the two eigenvalues of the covariance matrix *λ*_1_*, λ*_2_ which relate to individual variances along each principal axis. L) Average Mahalanobis distances for the three experiments across-days, shown for B1. Each dot represents an individual day, and the color-coded lines are linear regression fits for each experiment. Combined, CL1 and CL2 exhibited a significant linear increase in the mean Mahalanobis distances across days. M) Average Mahalanobis distances for the three experiments across-days, shown for B2. N-O) Boxplots of neural variances across all sessions and movements between the three experiments for B1 and B2 respectively.

Each day, we carried out the procedure of first starting with OL and initializing a new decoder, and culminating in CL2. Note that in these initial experiments, there was no robot motion, either during open-loop data collection or during closed-loop control; we were primarily focused on the stability and flexibility of the discrete neural commands in the open-loop and closed-loop settings.

For optimal somatotopy-based BCI decoding, neural features were preprocessed using spatial and temporal averaging and running the decoder at slower rates (between 5-10Hz, Methods and see (Branco et al., 2018)). These preprocessing steps significantly increased spatial SNR and improved single-trial consistency (see Fig. S3A-B for a single session, *p* ≤ 0.01).

### 2.3. Discriminability of representations is higher during closed-loop BCI control

Representative trial-level confusion matrices for three types of experiments (OL, CL1, and CL2) for a given day are shown in Fig. 2D-F for B1. The mean decoding accuracies across all sessions are shown in Fig. 2G for B1. While an offline decoder could classify held-out OL trials well above chance levels (68.45%, [95% C.I. 64.22 - 72.79], chance = 14.29%, *p* ≤ 0.01, Fig. 2G), decoding accuracies were comparatively and significantly greater during both closed-loop experiments (CL1: 80.13%, [74.34% - 85.38%], *t* (9) = 4.8*, p* = 9.668 × 10^−4^); CL2: 85.71% [80.12% - 90.71%], *t* (9) = 7.283*, p* = 4.646 × 10^−4^). In B2, while cross-validated open-loop decoding accuracies were at chance levels, decoding accuracies during closed-loop control were significantly above chance (Fig. S2B). Moreover, the percentage increase each day in decoding accuracies relative to the OL experiment was significantly higher in CL2 than CL1 across both subjects. This highlighted the efficacy of updating the decoder with closed-loop data (Jarosiewicz et al., 2013) (Fig. 2H, CL1 vs. CL2 relative to OL reference, black dotted line, *mixed effect model*, *t* (26) = 2.636*, p* = 0.0139). Notably, OL decoding performance in B1 did not improve across-days even with sustained practice with the BCI linear model intercept *β* = 0.012 ± 0.01*, t* (8) = 1.16*, p* = 0.277). Behavioral data thus indicated that sensorimotor representations were decoded better during closed-loop control especially after a decoder update, while they tended to remain stable in the open-loop setting. We then sought to understand how the open-loop neural representational structure adapted to the BCI context both within and across sessions.

### 2.4. Changes in representational statistics with closed-loop BCI

To visualize high-dimensional network-wide cortical activity and contrast the neural correlates of open-loop and closed-loop behavior, we performed dimensionality reduction to identify a neural manifold. This low-dimensional latent space, via a neural-network based autoencoder (Hinton and Salakhutdinov, 2006; Pandarinath et al., 2018; Pailla et al., 2019; Zhou and Wei, 2020; Talukder et al., 2022; Langdon et al., 2023)) Methods) succinctly captures high-dimensional motor control data for the repertoire across all three experiments. We built this manifold separately for each session with each day’s data. Neural activity of the first and last day captured on their respective manifolds are shown in Fig. 2I and 2J respectively for B1. Inspection of latent-space activity revealed immediate within-session shifts and separation between movements’ latent distributions when transitioning from open-loop to closed-loop control. There also appeared notable reductions in each movement’s neural variance specific to the BCI-context as open-loop neural variance appeared to be consistently larger. To quantify these effects, we measured the a) Mahalanobis distance, which measures the pairwise statistical separation between movements’ low-dimensional neural distributions (differences in means weighted by pooled variance), and b) the neural variance or spread of distributions in latent-space (Fig. 2K).

Confirming our observations, closed-loop control elicited greater average Mahalanobis distances as compared to open-loop data (Fig. 2L and 2M, main effect of experiment-type in B1 *rmANVOA F* (2, 18) = 145.052*, p* = 7.88 × 10^−15^, distances in B1, CL2: 45.06 ± 2.353 S.E., CL1: 33.2 ± 2.292 S.E., OL: 8.34 ± 0.446 S.E.; main effect of experiment-type in B2 *rmANVOA F* (2, 6) = 16.54*, p* = 0.0036, distances in B2, CL2: 0.554 ± 0.1 S.E., CL1: 0.26 ± 0.052 S.E., OL: 0.082 ± 0.12 S.E.). Notably, neural activity in CL2 elicited the highest average Mahalanobis distances in both subjects (Fig. 2L and 2M). With B1, in whom we could track latent space statistics across a longer period, the average Mahalanobis distances increased with practice across both closed-loop control experiments (Fig. 2L, *mixed-effect model intercept β* = 1.612 ± 0.614*, t* (18) = 2.623*, p* = 0.0172). Strikingly, the average separation between representational distributions remained stable in the open-loop setting (Fig. 2L, *β* = 0.2 ± 0.149*, t* (8) = 1.34*, p* = 0.216). We observed a similar trend of increasing separation across-sessions during CL2 in B2 albeit with a shorter number of days (Fig. 2M). Importantly, analyses of Mahalanobis distances in the original full high-dimensional feature space also matched our observations in latent-space (Fig. S4).

Analyses of neural variances revealed significant differences between the three experiments (Fig. 2N and Fig. 2O, B1: *mixed-effect t* (208) = 10.424*, p* = 9.694 × 10^−21^, B2: *mixed-effect t* (46) = 7.142*, p* = 5.586 × 10^−9^). Variance was consistently higher in the open-loop setting compared to closed-loop control (Fig. 2N and 2O) and was lowest in CL2 (two-sided *t-tests*, B1, OL vs. CL1: *t*(138) = 8.804, *p* = 4.898 × 10^−15^, OL vs. CL2: *t*(138) = 9.416, *p* = 1.452 × 10^−16^, CL2 vs. CL1: *t*(138) = 1.905, *p* = 0.0588; two-sided *t-tests* in B2, OL vs. CL1: *t*(30) = 4.78, *p* = 4.32 × 10^−5^, OL vs. CL2: *t*(30) = 7.257, *p* = 4.434 × 10^−8^, CL2 vs. CL1: *t*(30) = 2.612, *p* = 0.0139). On average, neural variance in OL was 4X higher than for CL in both B1 and B2. In contrast, the increase in just the distance between means during closed-loop control was only 1.65 times that of the open-loop setting.

This highlighted that changes in Mahalanobis distances were thus primarily driven by variance reductions. These main findings held even when expanding the number of actions (see Fig. S5 with a larger action set in B1).

The differences in latent-space variance between the three experiments notably did not involve variability in neural con-trol maps; rather these were reflective of modulations of neural variability at the channel-feature level. This is evidenced by in-variant cortical networks for the three experiments after passing latent-space activity through the decoder layers of the autoen-coder (Fig. 3). Specifically, we found highly preserved neural maps between OL, CL1, and CL2 each day across the three experiments and frequency bands for the movement repertoire (spatial *R*^2^ for all days, Fig. 3B and Fig. 3C). The average *R*^2^ across all three frequency bands was 0.972 [0.966 − 0.977] for B1 and 0.96 [0.956-0.972] for B2. Instead, we observed re-duced neural variability at the individual channel-feature level during CL1 and CL2. This is shown at two exemplar channels and movements in Fig. 3D for *γ_H_*. Across all three frequency bands and both subjects, closed-loop experiments were always characterized by significantly lower variance in channel activ-ity across the grid (in terms of channels’ standard deviation) as compared to open-loop experiments (Fig. 3E-F, *2-sample KS tests*, *p* ≤ 0.05). In addition, CL2 elicited lower variances across the grid than CL1 in *γ_H_* (CL2 vs. CL1 *2-sample KS statistic* = 0.165*, p* = 0.004) and *β* (CL2 vs. CL1 *2-sample KS statistic* = 0.13*, p* = 0.046) for B1 and in *δ* (CL2 vs. CL1 *2-sample KS statistic* = 0.36*, p* = 9.737 × 10^−8^) for B2.

**Figure 3:**
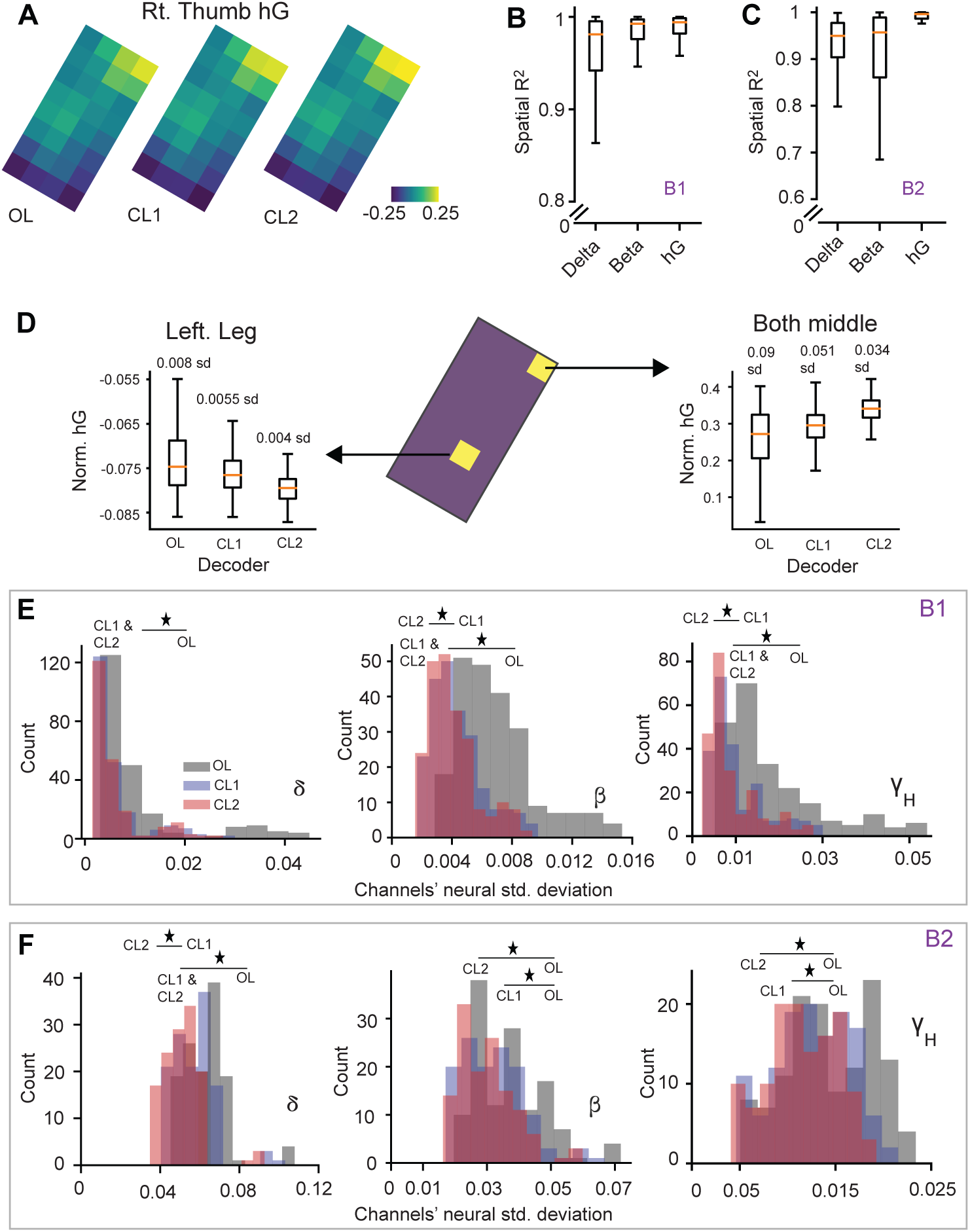
BCI-specific reductions in neural variance at the channel-feature level across the ECoG grid. A) Single-session average right thumb *γ_H_* activity across the grid for the three experiments. B-C) Boxplots of the pairwise correlation coefficient *R*^2^ between the average movement-specific neural maps of OL, CL1, and CL2 within each session and for all movement types. D) Example of BCI-specific within-session reductions in *γ_H_* neural variance (in terms of standard deviation) for two example movements, across two different channels on the grid. Boxplots correspond to data from the active portions of the task across all trials for the two specific channels. E-F) Histograms of the channels’ standard deviation in neural activity across the grid color-coded for OL, CL1, and CL2, and shown individually for the three neural features (*δ, β, γ_H_*) for B1 and B2. The star sign represents a significant difference between the distributions grouped by the horizontal line by a two-sided *Kolmogorov-Smirnov test*.

### 2.5. Stability of action representations across-days

We then sought to understand whether continued BCI practice across-days affected stability of the cortical networks governing movements i.e., the across-day stability of the manifold. Our representational analyses so far were all done within-session by building a manifold for each day. This limits understanding if the construct of action representations was preserved across-days. We thus examined the across-day manifold stability by comparing each session’s manifold to others. We specifically evaluated the similarity of individual layers between any two days’ autoencoders (Fig. 4A) using the Centered Kernal Alignment method or CKA ((Kornblith et al.), Methods). An example of the layer-wise similarities between two exemplar days’ autoencoders for B1 is shown in Fig. 4B (red vertical line) assessed against a null distribution (light blue histogram). It can be seen in Fig. 4B that the CKA similarities between the two days were highly significant at individual layers. The vast proportion of all pairwise compar-isons between days had almost all layers to be significantly similar (Fig. 4C, *p* = 0.05, *FDR* corrected). In B1, 100% of all pairwise comparisons had all 5 layers to be significantly similar to each other, and in B2, 83.33% of all pairwise com-parisons had all layers similar to each other. This suggested remarkable stability of well-rehearsed stereotyped movements, characterized by an across-day preserved manifold, even with novel and continued contextual adaptation to the BCI.

**Figure 4:**
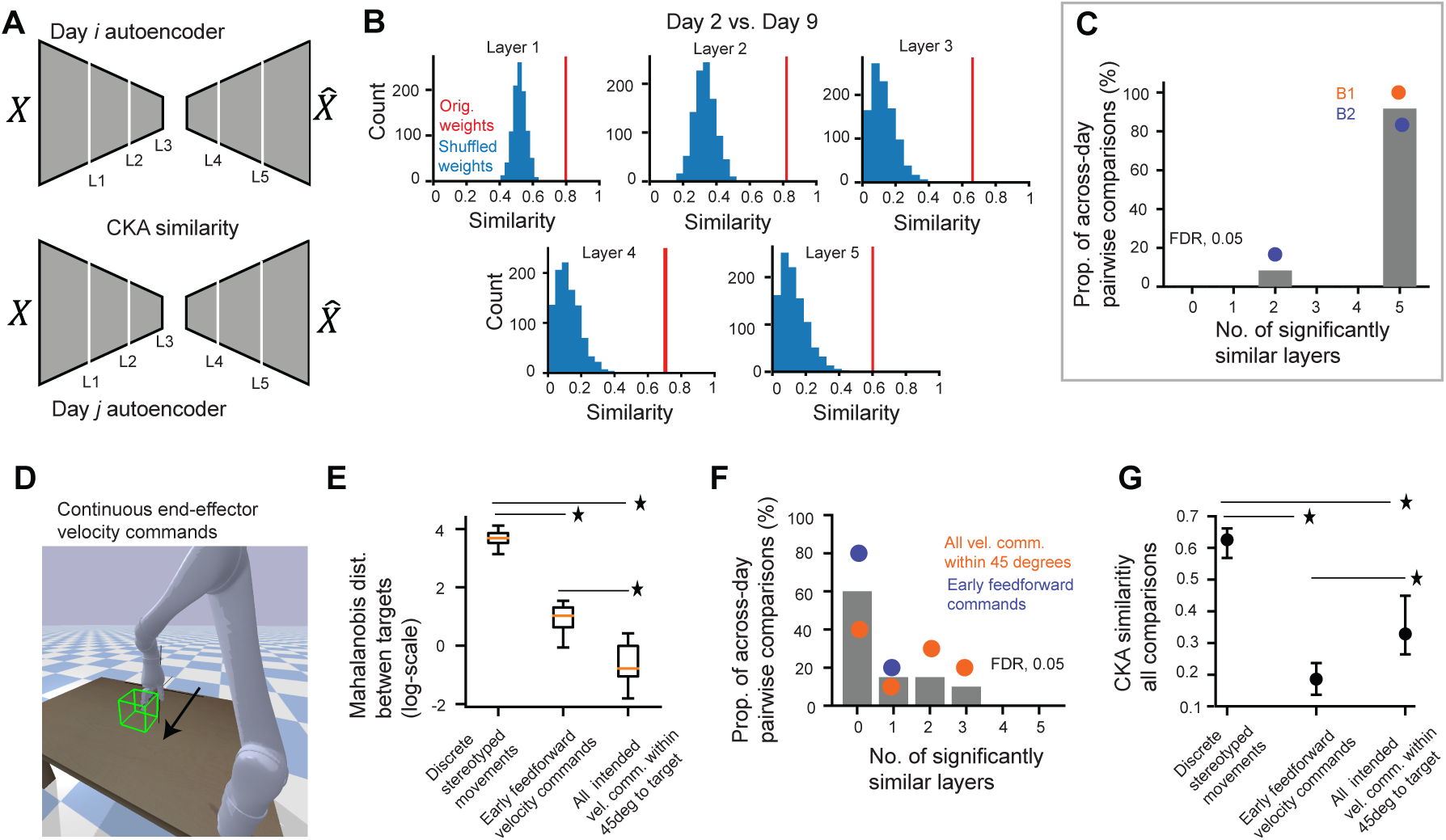
A preserved neural manifold characterizes the across-day representational stability of daily initialized discrete well-rehearsed movements unlike denovo continuous end effector motor commands. A) The CKA metric was used to compute the layer-wise similarity between any two days’ autoencoders. Similarity was assessed at five distinct hidden layers, including the bottle-neck capturing latent-space activations (Layer 3). B) Example CKA similarities between Day 2 and Day 9’s autoencoders at each of the five layers (red vertical line) against a null distribution of similarity values (blue histogram). All five layers were significantly similar to each other. C) The proportion of all across-day pairwise comparisons between the daily autoencoders representing stereotyped movements (Y-axis) that had a significantly similar number of layers (X-axis) to each other at the *p* = 0.05, (*FDR*) level. Each dot represents the proportions of a specific subject. D) Snapshot of center-out robot movement requiring imagined biomimetic end-point velocity commands. Collected neural and kinematic data were then used to seed and subsequently update a ReFit-Kalman Filter. E) Boxplots of the Mahalanobis distances between neural commands during closed-loop control to center-out axial directions collated across all sessions, either via discrete stereotyped representations or two types of continuous velocity commands. The star sign represents a significant difference in means between variables grouped by the horizontal line. F) The proportion of all across-day pairwise comparisons between the daily autoencoders representing denovo velocity commands (Y-axis) that had a significantly similar number of layers (X-axis) to each other at the *p* = 0.05, (*FDR*) level. Each dot represents a particular type of denovo velocity command in subject B1. G) CKA similarities across all pairwise and layer-wise comparisons for the three types of representations.

We then wondered if similar stability existed for traditional continuous 3D end-effector control. These are typically initialized using observed actions and then adapted over time to operate a motor neuroprosthesis. In B1, with whom we were able to continue the study for a longer period, we collected daily imagined end-point biomimetic velocity commands as the user watched the virtual robotic arm move in a center-out fashion in 3D space to six axial targets (Fig. 4D). The imagined commands were then used to seed an online velocity ReFit Kalman Filter (ReFit-KF (Gilja et al., 2012)). The parameters of the Kalman Filter were updated within-session after multiple blocks of real-time fixed control using SmoothBatch (Orsborn et al., 2012). This formed open-loop (OL) and closed-loop (CL) control blocks analogous to the prior discrete experiments. We initialized and ran a daily Kalman Filter for five experimental days. First, we found that intended velocity commands with the ReFit-KF exhibited much less target-specific discernability than those of well-rehearsed representations (Fig. 4E, Fig. S6). The averaged latent Mahalanobis distances between closed-loop neural representations to targets were much higher with discrete stereotyped movements compared to velocity commands (Fig. 4E). We found this to be the case whether it was with respect to early feedforward velocity commands (first ˜2.5s of trial-start, *two-sample t-test*, *t* (28) = 11.65*, p* = 2.94 × 10^−12^) or all intended velocity commands to the target within a 45 degree tolerance (over the entire trial duration, *two-sample t-test*, *t* (28) = 12.356*, p* = 7.44×10^−13^). Between the latter two, early feedforward commands contained significantly more discernability (Fig. 4E, paired t-test, *t* (9) = 4.514*, p* = 1.46 × 10^−3^).

Importantly, the manifold characterizing daily-initialized continuous velocity commands tended to exhibit much less representational stability (Fig. 4F, Fig. 4G). Pairwise comparisons revealed that the majority of across-day manifold comparisons had no significant layers between them (Fig. 4F, 60% of all comparisons were not significant). CKA values were also higher for simple imagined movements compared to continuous velocity commands (Fig. 4G, bootstrapped test, stereotyped movements vs. early feedforward *p* ≤ 1 × 10^−4^, stereotyped vs. all intended velocities within a 45 degree tolerance *p* = 2 × 10^−4^). Interestingly, although neural represen-tations during early feedforward velocity control carried more target-specific discernability, they tended to be much less stable across-days. On average only 0.2 layers between daily autoen-coders were significantly similar. However, velocity commands within a 45 degree tolerance had 1.3 significant layers on av-erage (Fig, 4F). This is further highlighted by the contrasting CKA values between the two (Fig. 4G, early feedforward vs. all intended velocities with a 45 deg tolerance *p* = 0.0148).

### 2.6. Using representational stability for plug-and-play (PnP) BCI control in the presence of distributional drift

Our data have established that well-rehearsed action representations are characterized by a preserved manifold and a stable open-loop representational structure that can be flexibly upregulated across BCI contexts. However, computation of this representational structure is based on relative distances between movements. CKA also mean-centers data prior to computing activation similarities. As such, these are independent of across-day distributional drifts (especially in centroids), and hence the absolute location of the manifold. Such drifts may be likely even for stable representations (Bishop et al., 2014), and would imply that the interplay between representational stability and plasticity occurs on a manifold whose location is drifting across-days. Large across-day distributional shifts also limits the feasibility of a decoder based on stable representations to generalize across-days i.e., be plug-and-play (PnP) without requiring repeated recalibration (Silversmith et al., 2021).

To understand the drift in neural distributions, we evaluated day-specific differences in latent-space. As an example, B1’s open-loop right thumb movements across three random days in a common latent-space are shown in Fig. 5A. Examination of data strikingly revealed little distributional overlap between each day’s recordings. This is further highlighted in Fig. 5B detailing appreciable across-day drift in the means of channel-level activity over an across-day preserved neural map (shown for *γ_H_* from the hand-knob region). In general, for both B1 and B2, the day of recording in latent-space was highly discernible (Fig. 5C and 5D, 90.96% for the 10 days in B1 and 86.51% for the 5 days in B2). In contrast, centering and demeaning each day’ data resulted in accuracies to be indistinguishable from chance (B1: 10.88%, *t* (9) = 0.867*, p* = 0.408, *B*2 : 20.2%*, t*(4) = 0.113*, p* = 0.915). This suggested that although there was comparative representational stability for the repertoire of stereotyped movements, there was an across-day drift in the centroids of each day’s neural distribution. This concept is detailed in Fig. 5E wherein the location of the preserved neural manifold is affected across-days while maintaining stability in the representational structure.

**Figure 5:**
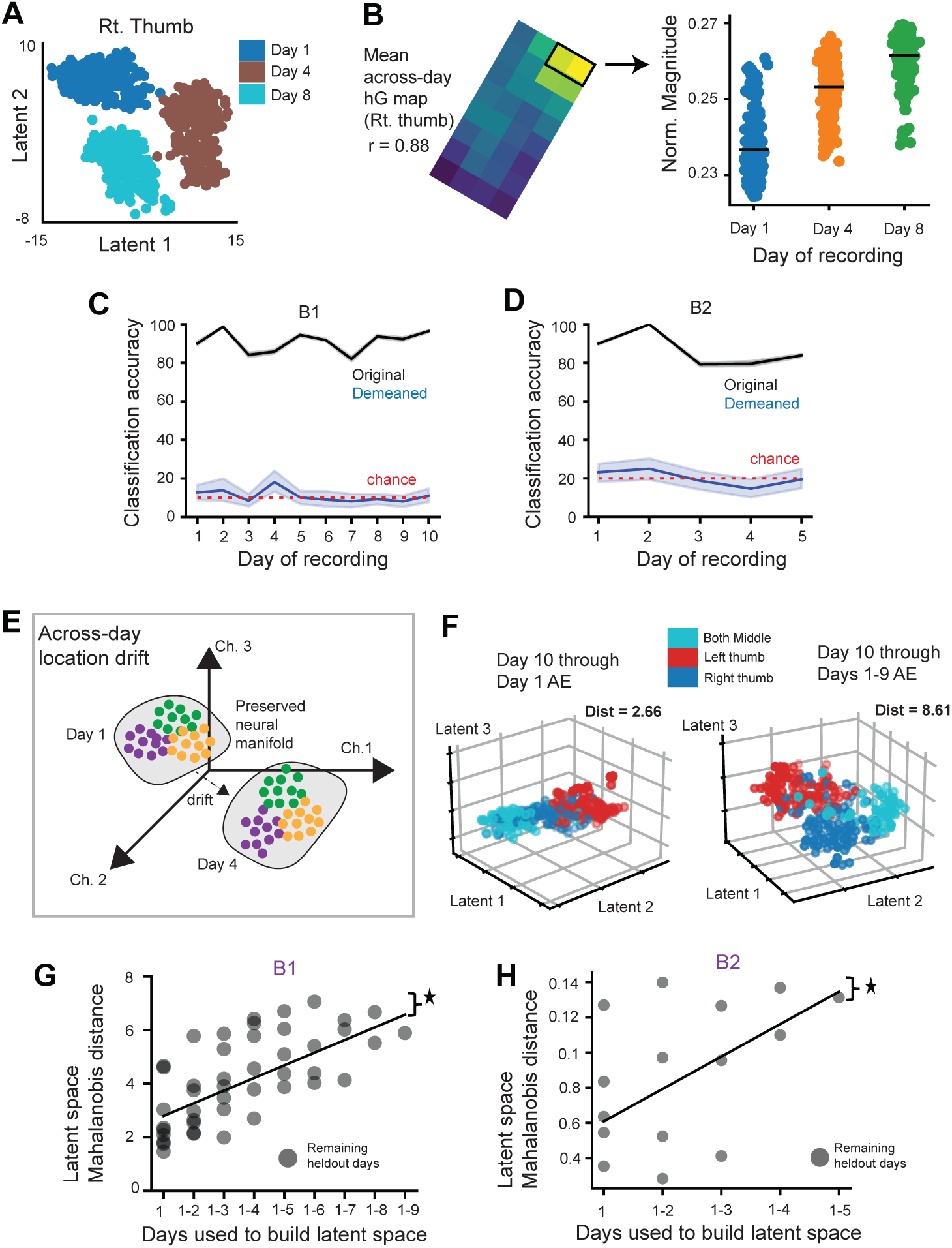
Using multi-day data to account for across-day drift in distributions of stable representations. A) Visualization of neural samples from three randomly chosen days of open-loop right thumb data in latent-space. B) Mean *γ_H_* neural map for the three random days (left, spatial *R* = 0.88), and shifts in the mean of M1/S1 channel-level activity (black rectangle) shown in distributions on the right. The filled dots represent neural samples color-coded by day, and the black vertical line depicts the mean. C-D) Cross-validated classification accuracies of discerning the day of recording (all movements) in latent-space for B1 and B2 when retaining the mean of each day’s data (black) or after mean-centering (blue). Chance is depicted by the red dotted line in both plots. E) The across-day drift in the neural mean activity affects the daily centroid of a stable neural manifold in high-dimensional channel-feature space. Each dot on the manifold is symbolic of neural data samples, color-coded by exemplar movement-types. Even though the representational structure i.e., relative locations of the movements on the preserved manifold is stable, the manifold location drifts across-days due to daily distributional shifts. This potentially affects the generalizability of a daily decoder. F) Projection of hand-specific movements (rt. thumb, lt. thumb, both middle fingers) from the 10^th^ experimental day onto a manifold built using only Day 1 data (left) or through a manifold built using multi-day data from Day 1 to 9 (right). Each filled circle represents a neural sample color-coded by movement-type. The average of pairwise Mahalanobis distances between the three movements in latent-space (Dist.) is detailed for each manifold. G-H) Mean of all pairwise Mahalanobis distances between all movements’ open-loop data (Y-axis) from held-out days when projected onto a manifold built with a cumulatively increasing number of days before held-out days (X-axis), for B1 and B2. Line represents the best linear regression fit to all data. Star signs indicate a significant slope to regression fit.

How might we account for this drift for a generalizable representations-based BCI decoder? Given the relative representational stability of simple actions and the flexibility in regulating their statistics, the user could overcome this shift in the distributional centroids via plasticity and successfully operate a daily decoder. Alternatively, using multi-day data to sample distribution shifts of stable representations (Rule et al., 2020; Willett et al., 2021) might enable PnP performance without solely having to rely on neuroplasticity. This can potentially speed up time to achieving long-term stable control. To test this hypothesis offline, we rebuilt the manifold using open-loop data from a cumulatively increasing number of days. We then evaluated how well it captured the discriminability between movements on held-out days. As an example, we projected all of B1’s hand movements’ neural data from the 10^th^ day onto a manifold built using either the first day alone (Fig. 5F, left) or through the first 9 days (Fig. 5F, right). The manifold built using 9 days appeared to better capture separations between hand movements than a manifold built using only the first day.

In general, we found that the separation between movements at any given day linearly increased as a function of the number of prior days used to build the manifold. This effect can be seen in Fig. 5G and 5H (linear model slope on the median at each day, B1: *t* (7) = 6.38*, p* = 1.87 × 10^−4^; B2: *t* (3) = 10.3*, p* = 0.002). Thus, for generalizability and PnP control in the setting of stable representations, there appeared to be benefits in utilizing multi-day data to sample shifts in the centroids of neural activity. This also suggested that the drift was not completely stochastic and might be within plasticity bounds (Rule et al., 2020; Masset et al., 2022). We thus used all open-loop and closed-loop data from all experimental days to seed a PnP decoder (feedforward neural network, Methods), with the hypothesis that the resultant decoder would generalize well online across-days. All results henceforth pertain to B1 who was able to continue in our BCI sessions. We tested the effectiveness of the PnP decoder when the subject was instructed to select cued virtual targets as rapidly and accurately as possible in the same discrete experiment (detailed earlier in Fig. 2C).

Accuracies and the average time taken to select a target (both dependent on the accumulation of decoding probabilities exceeding a threshold, Methods) within experimental blocks across a 4-week PnP window are shown in Fig. 6A and 6B respectively. The average accuracy across the entire PnP period was 87.43% [84.0%-90.8%] and the average time to target was 1.195s [1.087s-1.33s]. This resulted in a mean bitrate of 1.72bps [1.51-1.94] (Fig. 6C), an information-theoretic measure dependent on both accuracies as well as the time taken (Nuyujukian et al., 2014). An exemplar high-accuracy block is shown in Supplementary Video 1. We then updated the weights of the decoder using the first experiment’s data and ran a second experiment across a longer PnP window (Fig. 6D-F). The average accuracy was 80.71% [77.6%-83.5%] and the average time to target was 0.875s [0.842s-0.917s] resulting in an effective mean bit rate of 1.848bps [1.68-2.02]. We thus largely replicated the stability and performance of the first PnP experiment. Note that the accuracies in the PnP experiments were similar to a daily decoder (detailed in Fig. 2). Across both experiments, the mean effective bit rate was 1.8bps [1.66-1.93] and the highest was 3.351bps, achieved just by using somatotopic spatial representations through a feedforward network.

**Figure 6:**
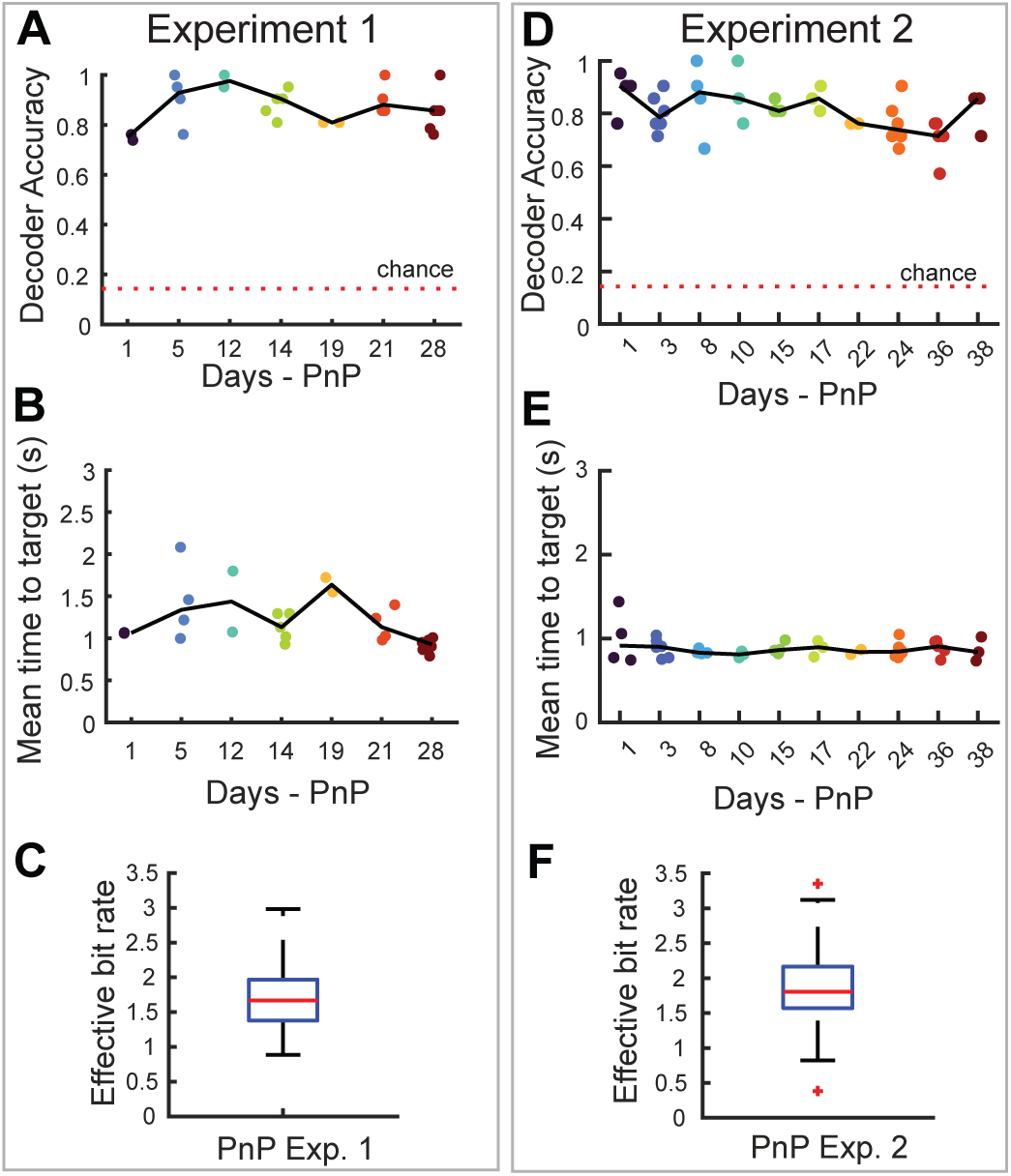
PnP decoder performance using discrete representations. A) Decoder accuracy in discerning the movement class in the first long-term PnP experiment. Each filled circle represents accuracy from a block of continuous trials, and the color-coding corresponds to the day of the PnP experiment. The line connects the median performance of each day. B) The mean time taken for the PnP decoder to arrive at a decision in the first long-term PnP experiment. Each filled circle represents a block of trials, and the line connects the median performance of each day. The color-coding corresponds to the day of the PnP experiment. C) Boxplots of the bitrates from individual blocks across all days of the first PnP experiment. D-F) Performance in terms of decoder accuracy, mean time to target and bitrates in the 2nd PnP experiment. In F) outliers are represented by the red squares.

### 2.7. Translating PnP decoder for continuous 3D end-point control of a virtual robotic arm

Having established that by accounting for drift, discrete action representations could be flexibly used as long-term stable BCI control signals, we then sought to translate our findings for continuous hDoF (high degree of freedom) PnP control. Within the same 3D virtual environment, we enabled continuous robot motion using a framework detailed in Fig. 7A. Specifically, the body-centric discrete decodes along the cartesian directions were now used as actual velocity impulses *u* (*t*), scaled by gain *B* to alter the continuous end-point 3D kinematics (state *X* (*t*) of the robot, Fig. 7A). Robot dynamics (*A* matrix, Fig. 7A) were modeled to follow first-order laws of motion with smoothly decaying velocities in the absence of neural drive. The discrete representation corresponding to the origin (‘both middle fingers’) was used as a ‘grasp’ command that froze the robot’s position and performed a ‘grasp’ if the hand were within an object’s virtual boundaries. We termed this control framework IBID for Input-Based Integrated control of continuous Dynamics (Fig. 7A). Note that the neuroprosthetic is continuous i.e., the robot’s position spans all 3D space with continuous first-order dynamics; it is the neural velocity input to the system that is discrete (at momentary intervals) when the user wants to engage the robot. This decoding framework allows for highly accurate 3D center-out trajectories along axial directions (Fig. S6, compared with a denovo ReFit-KF using ECoG signals). It also allows for “coasting” where the user can disengage and allow the robot’s internal dynamics to take over (Fig. S7A). The efficacy of IBID in precision trajectory control was then tested in a ‘path-tracing task’. This task required the user to maintain stereotyped representational states over long time-periods and follow a predetermined path (Fig. 7B, top) to grasp a virtual cube (Fig. 7B). Similar to the discrete experiment, we still provided feedback of the decoder output via a red arrow. An example high-performance video for this task is shown in Supplementary Video 2. We observed highly accurate path traces (single-session examples in Fig. 7C).

**Figure 7:**
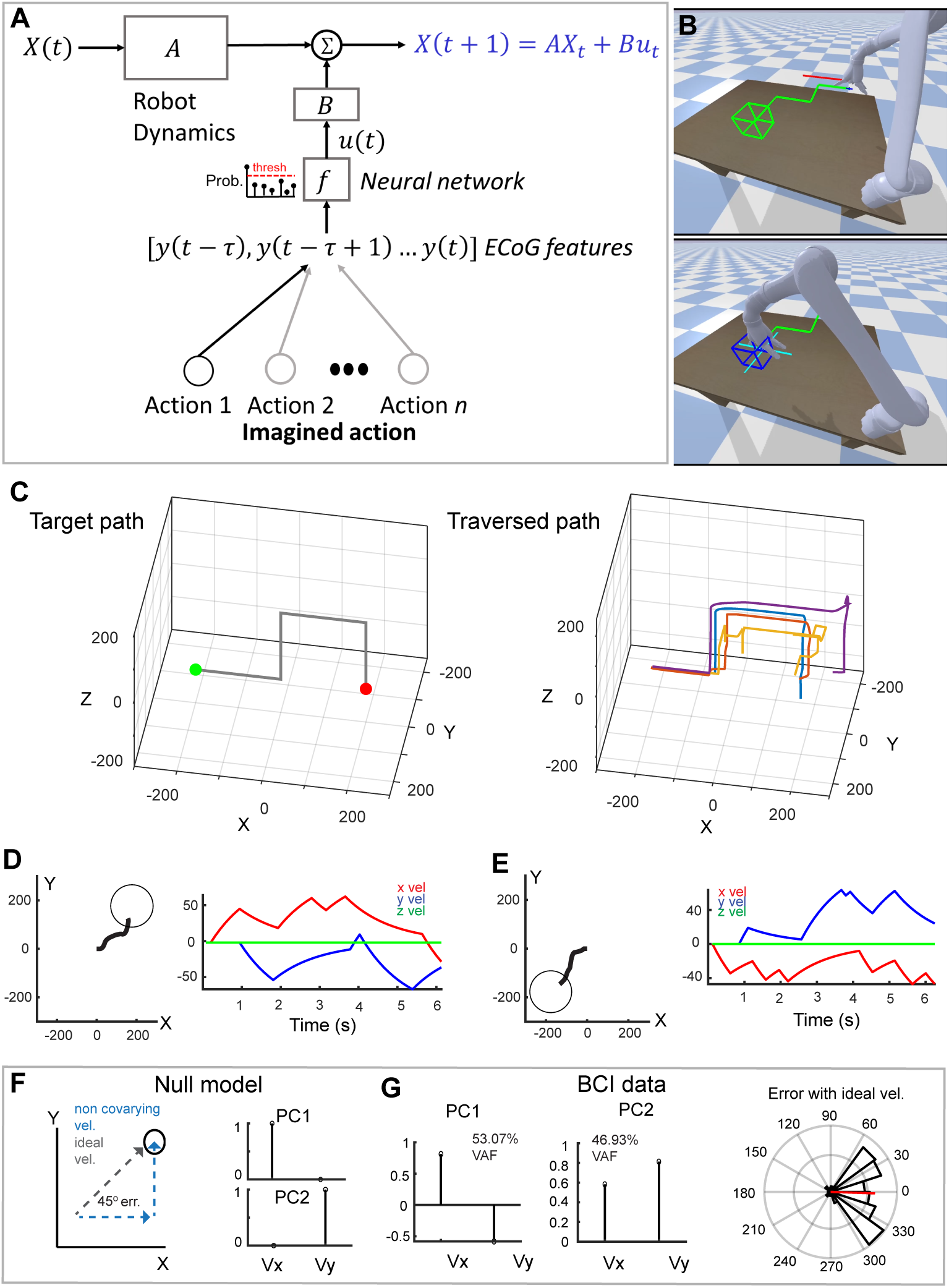
Input-based integrated control of continuous dynamics (IBID) using discrete representations. A) Algorithm used for continuous 3D end-point position control of a robotic arm using representations of discrete well-rehearsed actions as velocity inputs to alter ongoing first-order robot dynamics. B) (Top) Example of the path-tracing task that required the user to stay on the predetermined path (green line) and grasp the cube at the end of the path. The real-time instantaneous decoded direction from the PnP decoder along which velocity inputs are applied is shown by the red arrow. (Bottom) A grasp command changes the arrow to a stationary cross and the cube changes color when selected. C) Example of ground truth path the subject was asked to follow (left) and single trials (color-coded) examples of the subject accurately tracing the path within a session (right). D-E) Example trials (top view in 3D space) showing diagonal paths using the IBID framework when reaching a target along a 2D plane (left figure in both plots). The velocities driving the system is shown on the right in both plots. F) Null ‘etch-a-sketch’ model of non-integrated control wherein the user takes perpendicular steps along the axial directions to reach diagonal targets (left). This would always result in a 45-degree error between the ideal instantaneous velocity towards the target and recorded velocities. PCA on such kinematic data would then show no covariation i.e., the weights of the PCs would recapitulate the cardinal directions. G) (Left and middle) Weights of the two PCs from 2D velocity data during BCI control with IBID to reach diagonal targets as depicted in Fig. 7D and 7E. There was significant covariation in velocities. (Right) The polar histogram of the angular errors between instantaneous BCI velocities and the ideal velocities.

It should also be noted that our decoding framework does not preclude ‘off-axis’ control. Specifically, by serially switching between representational states more rapidly, diagonal trajectories can be executed, i.e., characterized by coarticulation of multiple axial velocities. Example off-axis trajectories with precise coarticulate velocity control in 3D space are shown in Fig. 7D and 7E. Such continuous off-axis trajectories are distinct from non-integrated perpendicular steps to reach diagonal targets (‘etch-a-sketch’ control). The null model for this latter scenario is shown in Fig. 7F for exemplar 2D velocity control in 3D space, characterized by directional non-integrated steps only along the cardinal axes. Non-integrated steps result in a constant error of 45 degrees with the direction of the ideal velocity straight towards the target (Fig. 7F). The lack of co-variation also implies that principal component analyses (PCA) on kinematic data would recapitulate the cardinal directions as PCs (Fig. 7F). In contrast, data from B1 revealed significant covariation in velocities due to the integration in IBID (PCA, Fig. 7G left). In addition, the integrated IBID velocities on average were always in line with the ideal direction towards the target (Fig. 7G, right, error in difference between ideal and IBID: 1.885 degrees [95% C.I. -5.21degrees to 1.44degrees] *Rayleigh test-statistic* = 440.126*, p* = 4.81 × 10^−221^) unlike the prediction of the null model (45 degrees). B1 was also able to access off-axis directions by coactivating multiple stereotyped representations simultaneously e.g., imagining the right thumb plus head to access targets in the positive XY quadrant (Fig. S7B), further highlighting the flexibility of our decoding framework.

Finally, unlike approaches that have an explicit relationship between neural activity and end-effector kinematics (e.g., regression-based approaches) our framework allows for broader generalizability. For instance, we found that B1 could immediately and proficiently use the PnP decoder even with a significant perturbation to the visual perspective and without any recalibration or relearning control. Single-session confusion matrices when operating the PnP decoder from either an allocentric frame (86% accuracy) or a left-handed framework of reference (94% accuracy) are shown in Fig. S7C and S7D respectively. This further highlighted the remarkable flexibility of discrete representations as stable BCI control signals.

### 2.8. hDoF virtual control of reaching and grasping

We then proceeded to extend the IBID control framework to more complex hand-object interactions. Specifically, instead of only focusing on transport, we also assessed the control of a gripper. A common method to enable such hDoF control is via a mode-switch command (Jain and Argall) wherein the user can selectively transition between and operate within the two phases (Fig. 8A and 8B). The main impetus for this was to further assess the flexibility of representations in allowing changes in mapping from transport control to gripper control. For end-point reach control, we used the same IBID transport framework as described in Fig. 7A. However, the ‘both middle fingers’ action was now used as a mode-switch to toggle between the transport and gripper phases.

**Figure 8:**
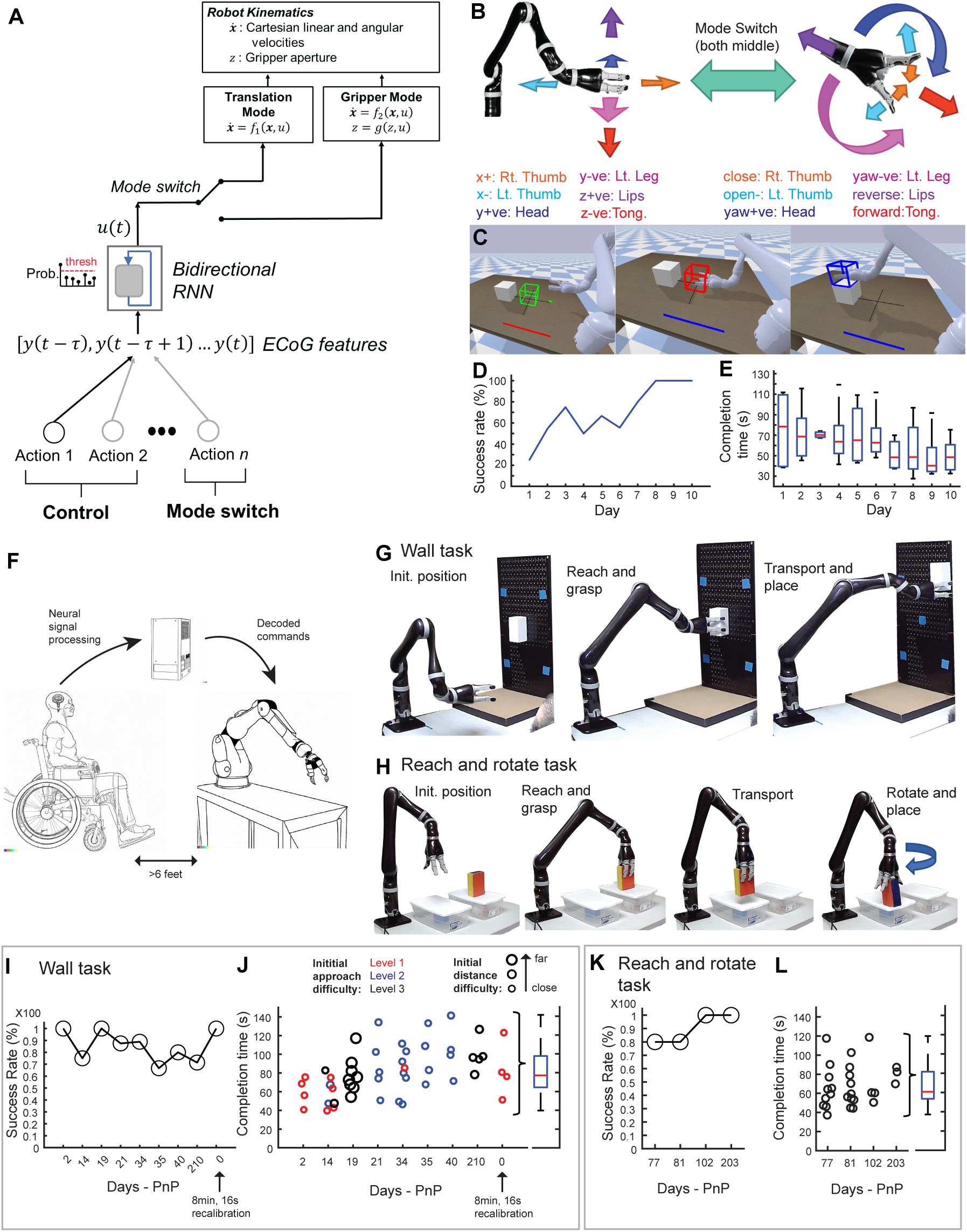
Long-term stable hDoF neuroprosthesis for complex reach-to-grasp and object manipulation tasks with the Kinova Jaco robot and using deep learning. A) Extension of the IBID framework to include control of gripper dynamics using a mode switch action. The feedforward PnP decoder was replaced by a PnP recurrent neural network when interfacing with a real-world Kinova Jaco robot. B) Overview of how discrete sensorimotor representations are flexibly reused and mapped for both gripper (right) and robot transport (left) dynamics via the mode switch action for the framework in Fig. 8A. Specifically, actions were remapped onto specific degrees of freedom (either gripper or end-point transport) and used via the mode switch. C) Snapshots of B1 performing a reach-to-grasp and transport task in the virtual environment using the framework detailed in Fig. 8A. The task required B1 to reach, grasp and place the cube onto the white box. The horizontal line at the bottom of the workspace table changes color to denote the mode-switch operation (orange – transport mode, blue – gripper mode). D) Success rates and E) task completion times across-days in the reach to grasp and transport tasks with the mode-switch in the virtual environment, highlighting increased proficiency with practice. F) Cartoon showing the physical setup of B1 with respect to the Kinova Jaco robotic arm. G) Snapshots of the wall task which required B1 to reach and grasp an object and then transport and place it along one of four cued corners of the wall. H) Snapshots of the lateral rotate task which required B1 to reach and grasp an object, transport and rotate it before placing it on a target box. I) Success rate each experimental day in successfully performing the wall task for all target difficulties. Days, since the weights of the RNN were held fixed (plug-and-play, PnP), are shown in the X-axis. Recalibrating the decoder resets the days back to zero. J) Task completion times each experimental day in the wall task. Each circle represents a trial, its size represents the initial distance between the Jaco and the target object, and its color represents the level of difficulty in the approach of the Jaco to the object. Days, since the weights of the RNN were held fixed (PnP), are shown in the X-axis. Recalibrating the decoder resets the days back to zero. K) Success rate each experimental day in successfully performing the lateral reach and rotate task. L) Task completion times each experimental day in the lateral reach and rotate task. Each circle represents a trial

During the gripper phase, the six actions used for cartesian reach control were remapped to gripper degrees-of-freedom (Fig. 8A and 8B). This included both rotational and transport dynamics i.e., rotation of the hand around a fixed axis in line with the gripper, and forward and backward translation of the hand to reach and withdraw from an object (Fig. 8B). Similar to the robot end-point’s dynamics, the gripper dynamics were also first-order and smoothly decayed in the absence of neural drive. Finally, two separate actions were used for opening and closing the gripper. While the flexibility of our framework allows for arbitrary choices in remapping, we sought to keep control as intuitive as possible, for example, by using hand-specific actions for gripper opening and closing. An example trial in the virtual environment of B1’s complex hDoF control using the mode switch is shown in Fig. 8C. The task required B1 to grasp the green cube, transport it and place it on top of the white box (Supplementary Video 3). Snapshots of B1’s performance during the trial are shown in Fig. 8C, and the sequence of neural commands discerned by the PnP decoder are shown in Fig. S8. During object interactions (grasp and drop), B1 successfully mode switched from transport to gripper mode and vice versa for the transport phase. In addition, the neural drive was not constant but momentary, wherein B1 selectively engaged control in a very precise manner (Fig. S8). B1 gained rapid proficiency in using the mode-switch with a few sessions of practice and was able to execute this task with 100% accuracy in roughly a week of practice (Fig. 8D) with a median task completion time of 45s (Fig. 8E).

### 2.9. Long-term control of grasping and object manipulation using the Kinova Jaco robot via deep learning

Having become comfortable with using the mode-switch for complex hDoF control, we then translated our findings and framework for a real-world hDoF neuroprosthesis with a physical version of the Jaco robotic arm (Fig. 8F). In this setting, we replaced the feedforward PnP decoder with a recurrent neural network - RNN (implemented as stacked biLSTMs, Methods (Hochreiter and Schmidhuber, 1997)). While a simpler decoder based on just spatial somatotopic representations can be easily trained and used, we found that a more complex decoder (Fig. S9) trained on longitudinal spatiotemporal ECoG data significantly boosted performance while offering similar PnP stability. Due to FDA regulations, the arm could not be placed close to B1, or wheelchair mounted and instead, had to be placed at a minimum distance of six feet from B1 on a table (Fig. 8F). Despite this suboptimal configuration, it did not impede B1’s ability to control the arm and gripper in two complex reach-to-grasp and object manipulation tasks that were designed to test performance with long-term PnP: a wall task (Fig. 8G) and a reach and rotation task (Fig. 8H) with varying levels of difficulty (Methods). For both tasks, the weights of the RNN were kept fixed and we tested long-term PnP performance.

B1 performed these tasks under full volitional control, i.e., without additional assistance. Performance in the wall task is shown in Fig. 8I and 8J and a demo is shown in Supplementary Video 4. The median success rate was 87.5% across all levels of difficulty, and the median time to complete the task in successful trials was 77.19s [72.07s-84.22s]. Notably, the PnP RNN could be stably used for nearly 7 months without recalibration, further suggestive of the fact that drift was within plasticity bounds. However, performance was higher in the first month both in terms of time to task completion (first month: 68.52s [56.12s-75.52s], all other days: 94.64s [78.26s – 103.13s]) and success rate (first month: 93.75%, all other days: 75.7%). This suggested that updates to the decoder might boost performance. For example, fine-tuning the PnP RNN immediately improved the success rate to 100% with a median time of 78.427s to complete trials (Fig. 8I and 8J). Notably, such fine-tuning required only 8min of calibration data with ground truth labels (Methods), highlighting the robustness of using simple action representations both for long-term control and for updates to the decoder in the presence of drift. PnP Performance in the reach and rotation task is shown in Fig. 8K and 8L, with a median success rate of 90% and median task completion times of 60.8s [54.76s-76.95s]. Notably, PnP performance in this task was very stable, even though we started this task approximately 2.5 months after fixing the weights of the PnP RNN (demo in Supplementary Video 4).

## 3. Discussion

Our results shed new light on how the human brain can maintain a stable representational structure on a preserved manifold for well-rehearsed imagined actions in the presence of drift, while at the same time flexibly adapting representational statistics in novel contexts. Importantly, in the absence of any feedback, there was a relatively high variance in the measured mesoscale responses for each imagined action. Accounting for distributional drift was key for enabling BCI control that generalized across days. This allowed our subjects to capitalize on practice from previous days to improve neuroprosthetic control. Overall, our study highlights the interplay between representational stability and plasticity and provides a framework that translates these findings for clinically relevant stable long-term neuroprosthetic control using mesoscale ECoG.

### 3.1. Mesoscale representational stability and plasticity in the presence of drift

The drift that we observed was across days and did not appear to be present in any given daily session. This suggests that it might be related to processes that modify neural representations over a day. One possibility is that our subjects slightly varied how they performed the imagined actions; we tried to account for this by ensuring that our instructions were highly specific and repeated frequently. We also used the same visual paradigm day-to-day to help constrain the ‘imagined’ actions. It is also unlikely that the drift could be due to pure stochasticity or unconstrained random walk outside bounds that could plausibly be overcome by plasticity (Rule et al., 2020; Masset et al., 2022). If this were the case, then PnP performance could be highly variable and might likely require a significantly larger number of across-day data. Instead, our analyses provided evidence of the efficacy of using multi-day; a PnP decoder built by sampling a limited number of days generalized remarkably well long-term. This suggests a latent meta-structure wherein the manifold’s drift itself might be constrained to another stable lower dimensional regime, which can be approximated by sampling distributional shifts across a limited number of days. Slight drift for well-rehearsed movements might provide some measure of constrained redundancy and tolerance in the neural code. This may allow for a larger neural space to exist for flexible activity patterns in a stable network of cortical neurons (Mau et al., 2020). This could also tie in with recent hypotheses that posit that it might be more advantageous for generalizability if network-level activity patterns were to drift time-varyingly around a stable attractor state than maintain a fixed-point (Driscoll et al., 2022). Nonetheless, accounting for this drift was extremely important for reliable decoding across days; without it, we did not observe ready across-day generalization. Without directly accounting for drift, PnP control likely requires long-term closed-loop decoder adaptation together with neural plasticity (Silversmith et al., 2021; Orsborn et al., 2014); this is likely to be more cumbersome and requires heavy user engagement, especially with increasing degrees of freedom.

Our data highlighted that even with drift and contextual adaptation to the BCI, the relative distances between movements on any given day during open-loop control were remarkably well preserved. In addition, the construct of the representations themselves was highly stable, characterized by a preserved neural manifold and somatotopy across both open-loop and closed-loop control. Prior fMRI neuroimaging work has suggested that cortical maps underlying well-learned actions in M1, such as finger movements, tend to remain stable when measured before and after motor learning (Beukema et al., 2019; Huang et al., 2013) as long there were no sensory perturbations of the end-effector (Kieliba et al., 2021). The increased resolution afforded by ECoG (including the ability to measure single-trial responses), along with the precise feedback-driven experimental manipulation of representations with long-term BCI, corroborate these findings and shed further light on representational statistics both in single trials during the task and across-days.

### 3.2. Comparison of mesoscale representations with single and multi-unit activity

Our long-term data also highlights the significant stability in mesoscale cortical representations of stereotyped movements, with a PnP decoder that could be stably used for many months. In contrast, prior single and multi-unit spiking data in cortical layers appears to show much less representational stability for highly stereotyped behaviors (Clopath et al., 2017; Rule et al., 2019; Schoonover et al., 2021). Unlike spike-based recordings, a single mesoscale ECoG channel represents the summed activity of a large population of neurons (Chang, 2015) and is therefore likely impervious to individual neurons dropping in and out and having instability in tuning properties to movement. In particular, population-level stability has been exploited for across-day generalizable spike-based BCI decoders even with high neuronal turnover rates (Degenhart et al., 2020; Karpowicz et al., 2022; Bashford et al., 2023). This requires a procedure for remapping population dynamics across days. Similarly, the reliability of representations in ECoG could have been leveraged to stabilize the decoder by recalibrating any given day’s recordings and decoder to a reference recording day. However, this reliance on recalibration would not be true PnP and instead, we found that by just sampling the across-day shifts in neural activity, we could build a PnP decoder that was stable for a long time-period.

### 3.3. Neurofeedback

A second main finding when interrogating the plasticity of well-defined, discrete movements is that the statistics of neural activity, both within each action and across the repertoire, were selectively adapted to the BCI with visual feedback. At least for the simpler discrete BCI task, our task design during closed-loop control primarily provided direct visual feedback about the decoded state i.e., the imagined action. This allowed for rapid learning within a session, with improved discriminability and performance gains relative to open-loop control. Such changes in the representational structure were primarily driven by variance reductions. This is consistent with past notions of ‘neurofeedback’, where abstracted feedback of neural activity patterns resulted in the modification of activity patterns in a single session (Fetz, 1969) and could be stabilized with practice across sessions (Ganguly and Carmena, 2009; Ganguly et al., 2011). Our results thus highlight that real-time feedback of pre-existing stable representations can immediately ‘quench’ neural variance and increase neural precision, thereby allowing the user to upregulate the perceptibility of sensorimotor representations. Indeed, the ability to reduce neural variance, especially at fast time scales, has been shown to be vital for perceptual decision-making (Arazi et al., 2017). Research has also found that reductions in neural variance are a potential mechanism used by the brain to stabilize neural activity around an attractor state when transitioning to task-evoked activity (Churchland et al., 2010; Ito et al., 2020). Importantly, we found that such variance regulation increased after a decoder update. For instance, it might have been the case that the improvements in CL2 decoding accuracies were driven purely by a decoder more sensitive to closed-loop data and with refined decision boundaries, without additional plasticity requirements. Instead, our results show that feedback of a decoder more aligned with the user’s closed-loop control state (Orsborn et al., 2014; Gilja et al., 2012; Silversmith et al., 2021; Jarosiewicz et al., 2013; Willsey et al., 2022) further allows the quenching of neural variance and upregulation of the representational structure.

It is also likely that as the user progressed towards more complex hDoF neuroprosthetic control, there were additional plasticity mechanisms recruited that were dependent on the feed-back of reaching and grasping stages. For example, for precise and complex robotic arm and hand control along with object-specific interactions, the user had to learn to stitch together precise sequences of actions (Fig. 8B-E). This may likely be related to neural plasticity which allows allow reliable and rapid transitions between ‘submovements’ (Lemke et al., 2019). Importantly, such complex control was achieved purely by integrating visual feedback with ongoing neural commands; here it might have helped that our decoder was running at a slower rate than typically used during continuous control, allowing visual feedback to be integrated. It is likely that incorporating additional sources of feedback, such as somatosensory inputs tied to neuroprosthetic control (Flesher et al., 2021; O’Doherty et al., 2011), can potentially increase the speed of complex continuous control.

### 3.4. Relevance to BCIs and hDoF motor neuroprostheses

From a translational perspective, we were able to achieve long-term complex neuroprosthetic control for reach-to-grasp and object manipulations with a fixed decoder, while also demonstrating performance comparable to spike-based recordings (Collinger et al., 2013; Wodlinger et al., 2014; Hochberg et al., 2012). Notably, the neuroprosthetic was completely under the user’s entire volitional control with no machine assistance. Our results thus present the lower bound of what may be possible using our hDoF decoding framework. The hDoF framework and PnP decoder’s performance also exceeded our prior work with 2D cursor control with a substantially shorter time to achieve PnP (Silversmith et al., 2021), further highlighting the benefits of leveraging existing stable represen-tations (Degenhart et al., 2018).

There is likely a continuum between schemes that facilitate control with existing neural representations versus those that heavily rely on plasticity; this distinction also becomes most clear when considering control schemes that generalize across days. For example, there is growing evidence that decoders that were initialized from natural movement control (or imagined biomimetic movements) do not readily generalize across days (Silversmith et al., 2021; Orsborn et al., 2014; Ganguly and Carmena, 2009; Ganguly et al., 2011; Athalye et al., 2017). In such studies, although the BCI decoder is seeded using existing representations, such as open-loop imagined arm movements, a continuous decoder places an additional transformation (either linear or nonlinear) on neural activity to decode instantaneous kinematics. During closed-loop real-time control, the user faces the burden of learning this transformation to successfully operate the BCI (Taylor et al., 2002; Carmena et al., 2003) leading to a two-learner system (Silversmith et al., 2021; Orsborn et al., 2014). Such an approach results in a neural map fundamentally different and sometimes completely orthogonal to the original representation used to seed the decoder (Silver-smith et al., 2021; Orsborn et al., 2014; Wander et al., 2013; Ganguly and Carmena, 2009; Athalye et al., 2017; Gulati et al., 2017). Indeed, we found that a ReFit-Kalman Filter initialized using visual observation resulted in poor across-day representational stability with much less target-specific discernability. In contrast, a discrete decoding framework perhaps does not incorporate the constraint of learning a transformation. As such, it seems to only reinforce the precision of existing stable representations, with visual feedback, to effectively learn proficient closed-loop control (Pancholi et al., 2023).

Our framework also highlights what type of commands might be best decodable for BCIs using mesoscale ECoG in paralyzed subjects (Chao et al., 2010; Ganguly et al., 2009; Schalk et al., 2007; Natraj et al., 2022; Leuthardt et al., 2004). Even years after injury, there appears to be a general preservation of cortical somatotopy and motor representations (Andersen and Aflalo, 2022). Its overall stability also offers the promise of scaling up to higher DoF (Gordon et al., 2023; Degenhart et al., 2018). Long-term stability and reliable performance is particularly important for hDoF neuroprosthetics. Especially given the complexity of hDOF BCIs, instability might necessitate long learning periods or excessive decoder recalibration. This can be time-consuming and hinder real-world adoption (Phillips and Zhao, 1993; Cordella et al., 2016). By identifying and using neural commands that are stable and whose drifts can be readily accounted for, we achieved long-term hDOF PnP control that can eventually improve real-world functionality in those with paralysis.

## 4. Contributions

N.N. designed the study, designed and implemented the real-time BCI signal processing system, decoders, and the hDoF control framework, performed experiments, analyzed data, and drafted the manuscript. S.S. implemented the real-time Kinova Jaco in virtual and real-world environments, the hDoF control framework, performed experiments and aided in drafting the manuscript. R.A. performed the initial calibration of the Kinova Jaco arm and hand, performed experiments, and aided in real-time system design. H.Y. and Y.G. aided in data collection. A.T.C. conducted patient recruitment and patient care. E.F.C. performed the surgical implantation of the ECoG array and performed patient care. K.G. conceived and supervised all aspects of the study. All authors read and revised the manuscript.

## 5. Acknowledgement

This work was funded by the National Institutes of Health through the NIH Director’s New Innovator Award Program, grant number [1 DP2 HD087955]. This work was also supported by the Weill Institute for Neurosciences at UCSF.

## 6. Methods

### 6.1. Clinical trial overview and participants

This study was approved by the US Food and Drug Administration (FDA) under an investigational device exemption (IDE) as part of the BCI for Restoration of Arm and Voice (BRAVO) clinical trial (NCT03698149). The IDE was specifically for the neural implant used in this study. Approval of the research paradigm and study protocol was obtained by the Committee on Human Research and Institutional Review Boards at the University of California, San Francisco. Participant B1 is a 41 year old right-handed male with extensive bilateral pontine strokes resulting in severe spastic tetraparesis and anarthria (Silver-smith et al., 2021; Moses et al., 2021; Metzger et al., 2022) and is wheel-chair bound with little to none upper limb movements (Medical Research Council scale of the right upper extremity: 1/5 for finger flexion, 3/5 for elbow flexion, 2/5 for elbow extension and 0/5 for other upper limb muscles; upper extremity Fugl–Meyer score of 7/66) and complete lower-limb paralysis. B1 was also a participant in our prior motor BCI study where we quantified his overt movement disabilities via video analy-sis (Silversmith et al., 2021). Participant B2 was a 56 year old male with end-stage ALS with complete paralysis from whom we had a limited window to collect data.

### 6.2. ECoG array and implantation

In both subjects, the neural implant was a 128-channel ECoG array in an 8X16 grid as shown in Fig. 1A. Total array dimensions were 6.7×3.2*cm*^2^, with a 2mm electrode recording surface contact and 4mm spacing between electrodes. The ECoG array is commercially available (PMT) and ECoG channels were connected to a transdermal connector and bonded to a pedestal (Blackrock Microsystems). Informed consent, medical and other surgical screening procedures were conducted before surgery. During surgery, the array of ECoG channels was placed on the cortical surface over sensorimotor including primary sensory and motor cortex, and areas overlapping with dorsal and ventral premotor areas. The pedestal connected to the ECoG channels was attached to the skull. In addition to the ECoG channels, reference leads were placed before surgical closure. The pedestal allowed a readout of the signals from the implant via a detachable digital head-stage and cable (Neuroplex E, Blackrock Microsystems) that digitized and transmitted minimally processed neural activity to a digital hub.

### 6.3. Real-time data acquisition

Neural signal measurements from the ECoG array relative to the hard reference wire were digitized and transmitted from the digital hub to a Neuroport system (Blackrock Microsystems). Here, adaptive line noise cancellation and anti-aliasing filters (0.3-500Hz band pass filtered signals) were applied before streaming the neural signals at 1Khz to a separate real-time BCI data processing computer via an Ethernet connection. The settings for the Neuroport system were controlled via software (Central, version 7.0.4, Blackrock Microsystems). There were then a series of signal processing, real-time visualization, and decoding steps using machine learning models on the real-time BCI machine (Ubuntu version 18.04 LTS) that were performed using custom code in MATLAB (Mathworks Inc.), Psychtoolbox, and in Python using NumPy, PyTorch, SciPy, scikit-learn, PyGame and PyBullet packages, with UDP communication between real-time BCI programs running in MATLAB and Python. As a re-referencing step to remove artifacts and background noise, we removed the common median signal across channels at each of the streaming 1KHz raw neural samples. Digital triggers from the real-time BCI machine tied to specific BCI task states and events were sent via an Arduino board to a separate data storage computer that also received raw streaming ECoG signals from the Neuroport system, allowing us to store task-synchronized raw neural data to disk.

### 6.4. Design of hDoF BCI and robot tasks

#### 6.4.1. Imagined movement data collection for somatotopy

To understand spatial somatotopy for a repertoire of imagined whole-body movements, we designed an event-related paradigm using Psychtoolbox. The structure of the trial is detailed in Fig. 1B. Each trial started with a text cue on the action to be imagined that was on screen for 4s, followed by the appearance of a red square indicating the subject to get ready (2s). The transitioning of the square color from red to green was the indicator for the subject to imagine performing the cued action (4s). After the trial ended, the screen turned black for a period of 3s as the inter-trial interval, and then the trial structure repeated. Each block of trials involved one repetition of the entire whole-body movement repertoire in a pseudorandom order, and we collected multiple blocks of trials across sessions. There were 30 such movements for B1 and a much limited number for B2 (∼10 movements that varied each session) given clinical constraints. There was no online control in this task and collected data with time-markers for the trial events were saved for offline analyses.

#### 6.4.2. Discrete movement decoding when mapped to cardinal 3D directions in the perspective of a virtual Jaco arm

The main experimental task to understand the regulation of the representational structure of stereotyped movements during hDoF control was performed in a custom Python environment with a virtual Kinova Jaco robotic arm (Kinova Robotics). The arm was placed in an egocentric right-handed perspective view of a workspace table. Each cardinal direction of 3D space, including the origin was mapped to a movement in a body-centric frame of reference that might be intuitive for control as shown in Fig. 2A. The X+ve and X-ve directions were mapped onto right and thumb movements, Y+ve and Y-ve movements were mapped onto head and left leg movements, Z +ve and Z-ve movements were mapped onto tongue and lip movements and the origin was mapped onto both middle fingers. Due to clinical and time constraints in B2, we were able to collect data from only four movements that were mapped to cardinal directions in a 2D workspace (X and Y axes). We first collected multiple trials of open-loop (OL) neural data during cued imagined access of targets (between 3-5s) along the cardinal directions. Each trial had fixed length inter-trial and preparatory intervals. Additionally, a consistent red arrow pointing in the direction of the cued movement was present on the screen for the entire trial duration, reinforcing the action to be imagined. In the case of the origin, a blue square at the origin was presented. Each block of trials had either 3 or 4 repetitions of each cardinal direction, resulting in either 21 or 28 trials per block for the seven movements. We collected multiple such blocks per session and resulting data were binned at the update rate of the decoder to be used in online closed-loop experiments. Closed-loop (CL) trials followed a similar structure to open-loop trials, with the major difference being that the subject now received visual feedback of their sensorimotor representation via the decoder output. Specifically, at each instant of the decoder update rate (between 5-10Hz) the direction of the red arrow was dependent on the classified movement. For example, if the right thumb was the decoded direction even if the cue was left leg, then the arrow pointed in the X+ve direction. In the case that both middle fingers were to be decoded then only a blue square at the origin was visually presented. Subjects were instructed to imagine the correct action so that the arrow continued to be pointing towards the cued target. In case the decoder probability of selecting an action did not exceed a user-defined threshold, then no visual output would be displayed to the user. The trial ended if the subject was able to maintain the arrow pointing toward the cued target for multiple consecutive bins (varying between 1-5 bins). There were two types of closed-loop blocks: CL1 involved a decoder seeded only on open-loop data and CL2 involved a decoder with both open-loop and CL1 data. To understand the stability and flexibility of representations, we conducted the experiment across multiple sessions, with each session starting with OL and culminating in CL2. This was done for 10 individual sessions with B1 and 4 sessions with B2 and 2 additional sessions with B2 involving only OL data. All online trials timed out after 5s.

Perturbations in perspective were introduced by either changing the handedness of the robot from right to left or the view, from egocentric to allocentric, while requiring the mapping between movements and intended axial directions to be consistent. In this manner, we were able to test whether the subject could flexibly and proficiently reuse stereotyped movement commands regardless of shifts in visual perspective. In addition, we also increased the number of mappings to 9 by including two additional actions, left and wrist movements, simulating the left and right rotation of the robotic arm in the virtual environment respectively. All the collected data with time-markers for the various task states and real-time decodes were saved for later offline analyses.

#### 6.4.3. Discrete PnP experiments

The plug-and-play (PnP) version of this experiment was conducted only with B1; the online task structure remained unchanged, but now the same PnP decoder was used every session. The instructions to the subject were to imagine the appropriate action and select the cued direction as accurately and quickly as possible. We applied a 1s-width mode filter to the classifier outputs (e.g., 5 bins when running at 5Hz) at the start of the active task portion to accumulate decisions when making a classification. Specifically, the mode of the decoded outputs within the running bin, including the null output when a decision probability does not exceed a threshold, was used to make a classification of the subject’s neural data. This places a lower bound of 600ms of correctly selecting the cued target when using a classifier running at 5Hz. All trials timed out after 5s.

#### 6.4.4. Continuous end-point kinematic control of the Jaco arm using discrete stereotyped movements as input velocity commands

To enable continuous end-point control of the Jaco arm, the discrete decodes were used as control inputs to a virtual dynamic model (damped mass-spring dynamics) of the robot. Each discrete decode was modeled as a direction-specific acceleration of fixed-magnitude (see Fig. 7A and Robot control frameworks in Methods). The action corresponding to the origin, ‘both middle fingers,’ was used as a stop command to freeze the robot, or as a grasp command to select objects if the end-point of the Jaco was within the boundaries of the object. We termed this framework as IBID or Input Based Integrated control of continuous Dynamics. In the virtual environment, we conducted a range of experiments with different objectives. First, to test center-out control, we collected trials requiring reaching and selecting target cubes placed along each axial direction, with the robot position resetting to the center at the end of the trial. In terms of task structure, the task states were similar to the discrete experiment, with fixed length inter-trial and preparatory intervals in addition to active control states. To test off-axis and coarticulate control, the targets were also placed at diagonal locations in the XY plane within the 3D environment. All online trials were set to time out within 15s. To further test the subjects’ proficiency in accurate velocity control, we designed a ‘path tracking’ task. This task required the subject to move the robot along a long predetermined path (colored in green, Fig. 7B) in 3D space and then ’grasp’ the cube target at the end of the path. Trials were set to time-out if the subject could not complete the task within 120s. In all the virtual experiments, the subject received two types of visual feedback: one was the actual motion of the robot itself in the virtual environment and the second was via the red arrow that pointed in the decoded direction (Fig. 7B). The subject thus received feedback on instantaneous velocity inputs in addition to estimating robot position, thereby allowing for more rapid error correction.

With the physical version of the Kinova Jaco, we also conducted center-out experiments to test continuous end-point control, with the robot placed at a distance greater than 6 feet in a left-handed perspective as shown in Fig. 8F due to FDA safety regulations. Before the start of each trial, the subject was verbally cued on which action to imagine and thus the direction to move the robot. The visual feedback received by the subject was only the physical movement of the robot itself. We also asked the subject to imagine the seventh action, ‘both mid-dle fingers’, in the real-world experiments; here the subject received visual feedback by an alternating color change (between red and blue) of a square on a computer screen that was placed 6 feet away and close to the physical robot.

#### 6.4.5. Continuous hDoF reach-to-grasp and object manipulation control with the Jaco in virtual and real-world environments

We extended our decoding framework using a mode-switch command to flexibly transition between end-point based transport phase and gripper-based control of the hand for grasping, releasing, rotating, and moving towards and away from objects (see Fig. 8A-B and Robot control frameworks in Methods). We then tested complex hDoF performance in a range of tasks in both virtual and real-world environments with the Kinova Jaco. The first task, in the virtual environment, was an object grasp and transport task (Fig. 8C). The robot was viewed from a right-handed perspective, and from the initial location, the subject was asked to reach toward a target cube placed on the center of a table in the virtual workspace. Having ‘grasped’ the object, the subject was then instructed to transport it without releasing or dropping it and place it on top of a white box. The color of the cube would change from green to red when grasped and would change to blue if released. The subject received three types of visual feedback: the robot’s position in the virtual environment, the decoded movement by the appearance and direction of a red arrow, and the mode-switch via a solid line at the bottom of the table. This line would change its color to indicate whether the subject was using the robot in the gripper (blue) or transport phase (red). The trial state structure was similar to other virtual tasks, with fixed length preparatory and inter-trial intervals and an active task phase. We collected multiple trials of this task over 10 experimental sessions to familiarize B1 with the mode-switch operation.

After the subject obtained sufficient practice and became familiar with the mode-switch operation, we transitioned to hDoF control of the physical Jaco arm in the real-world environment. We tested two complex hDoF tasks: a wall task and a top-down reach and rotation task, which required the subject to complete grasps from the side or top of an object, respectively. Specifically, the wall task required B1 to reach and grasp a rectangular object at the center of a wall, transport and place it on one of four targeted locations along the corners of the wall (Fig. 8G). The object had a magnet on one of its sides that was used to attach it to the wall. There were three levels of difficulties associated with the initial approach to the object: Level 1, requiring only a 1D reach to the object, for example, if the starting position of the gripper were to be horizontally in line with the object, Level 2, requiring B1 to navigate 2D to reach the object, for example, if the gripper were to be in the same 2D plane as the object, and Level 3, requiring B1 to navigate all 3D cartesian space to reach the object. Note that the robot was not clamped and all DoF were possible; decoding errors would therefore lose the advantage of having an easier path to the object. Placing the object on the wall required full transport and gripper control in 3D space; the subject had to withdraw the object from the wall and navigate to the target location, and then place the object on the target location and release the gripper. An additional layer of difficulty was a variable initial distance between the Jaco and the object. A trial was deemed to be successful if and only if B1 was able to complete all phases of the task successfully. The trial stop-time was recorded when B1 placed the object correctly at the target location and released the gripper. The lateral rotation task involved a reach and grasp of an object on a box, followed by transporting the object back to the starting location, and rotating and precisely placing the reoriented object at a target location (Fig. 8H). A trial was deemed to be accurate if and only if B1 was able to complete all phases of the task successfully and correctly orient the object when placing it on the target box. We collected multiple trials each session over many months and using a PnP recurrent neural network (RNN) based decoder. We initially focused experiments with the wall task given its complexity and introduced the rotation task much later on in the PnP experiments.

#### 6.4.6. Continuous end-point control of the virtual Jaco arm using ReFit Kalman Filter

To compare IBID end-point control with denovo biomimetic continuous velocity commands, we utilized a Re-Fit Kalman Filter whose parameters were updated using Smooth Batch (Silversmith et al., 2021; Gilja et al., 2012; Orsborn et al., 2012). We designed the experiments like the discrete action selection experiments. Each session started with open-loop trials where B1 observed the robot smoothly move from the initial starting location at the origin to targets along the axial directions in a center-out fashion. B1 was instructed to imagine biomimetic hand and arm commands to control the robot’s motion during observation. The imagined neural and generated kinematic data were then used to seed a ReFit Kalman Filter to decode instantaneous 3D end-point velocities. Online control was tested in a closed-loop control block (similar to CL1 in the discrete action selection experiment) where the ReFit-KF parameters were kept fixed. We then updated the ReFit-KF parameters using CL1 data and ran a second closed-loop control block with the updated decoder (similar to CL2 in the discrete action selection experiment). This process of seeding and updating a new ReFit-KF each day was repeated across five experimental sessions. Note that we did not carry over the weights across days, and reinitialized daily as we wanted to examine the daily stability of denovo end-point biomimetic velocity commands. Similar to other virtual experiments, each block had multiple trials and the trial structure was consistent, with a preparatory and inter-trial intervals and an active task state.

### 6.5. Real-time signal processing and feature extraction

#### 6.5.1. Spatial feature extraction for somatotopic decoders

For neural decoding using only spatial activity patterns, we computed the binned power of the streaming neural activity in three frequency bands at each channel of the 8X16 grid: *δ* (0.5- 4Hz), *β* (13-30Hz) and *γ_H_* (70-150Hz). The temporal width of the bins was dependent on the decoder update rate, which typically varied between 5 and 10Hz. For example, a decoder running at 5Hz would bin neural activity with 200ms intervals. Third-order Butterworth Infinite Impulse Response band-pass filters were used to extract the neural features real-time. To compute power more accurately in the slower *δ* band, the real-time PC had a running circular 2s buffer that was updated with raw neural activity at the decoder update rate. The filter was then applied to the entire 2s buffer and the most recent bin-width of *δ* band filtered activity was extracted. At the channel level, each sample of the filtered signal within the bin was squared and then averaged over time with a log transform to get an estimate of binned *δ* power. For faster rhythms, we utilized a filter bank approach before computing power and without the need for a running buffer. *β* power at each channel was computed by averaging binned power within 13-19 Hz, 19-30 Hz. *γ_H_* power at each channel was computed by averaging binned power within logarithmically spaced frequency bands 70-77 Hz, 77-85 Hz, 85-93 Hz, 93-102 Hz, 102-113 Hz, 113-124 Hz, 124-136 Hz, 136-150 Hz. To denoise signal features further and increase single-trial SNR and somatotopic differences given the geometry, density, and impedances of the ECoG grid, we performed both temporal and spatial averaging. The former was achieved by having a 1s online running average of the binned power within each of the three bands and at each channel. The latter was achieved via a 2 × 2 spatial averaging kernel that was rolled over the 8 × 16 grid with stride 2 (no padding) and within each frequency band. As a result, the 8 × 16 grid dimensions were reduced to 4 × 8 which were then flattened into a 32 × 1 vector for each of the three frequency bands. The concatenation of these vectors at each moment of the decoder update rate resulted in a 96 × 1 vector that formed the neural features. The final step was a *L*_2_ normalization of the 96-dimensional feature vector at each bin during the experiment. Instantaneous samples of the neural features were fed to neural networks for online decoding and to autoencoders offline for representational similarity analyses. At the start of each session, we collected two minutes of resting state data where the subject was instructed to relax and look at a blank screen. This formed the baseline data with respect to which raw experimental data were z-scored in real-time. Similarly, individual power at each channel within the three bands (before spatiotemporal smoothing) was also z-scored relative to their respective baseline power statistics.

#### 6.5.2. Spatiotemporal feature extraction for RNNs

For neural decoding using spatiotemporal activity patterns (e.g., with recurrent neural networks, RNNs), we extracted features differently. First, the real-time BCI system always had a running circular buffer that was typically between 800 and 1000ms. The buffer was updated with the most recent broad-band raw 1KHz streaming data at the update rate of the decoder. The raw data were z-scored to the baseline statistics at each channel (baseline mean and standard deviation), obtained via two minutes of resting state data at the start of each session. Note that the baseline z-scoring for the RNN was only on the raw streaming signals and not on the individual frequency features. Next, rather than have separate *δ* and *β*, we extracted a more general time-varying low-frequency signal via a 4th order low-pass Butterworth filter with a cutoff of 25 Hz. We retained the same logarithmically spaced filter bank and averaged over the sub-bands to compute time-varying *γ_H_* power at each chan-nel from the buffer. Each spatiotemporal feature data consisted of a matrix of dimensions 128×*t, t* ∈ (800, 100) which was then down-sampled in time by a factor of 10 using an anti-aliasing low-pass filter. Signals were then artifact corrected, wherein spikes in activity at any time and channel that was higher than a specified threshold would be replaced with extremely low variance Gaussian noise. Note that the down-sampled and hence smoothed low-frequency signal is analogous to low-frequency local motor potentials which have been shown to carry considerable movement-specific information in ECoG during both overt and BCI commands (Pistohl et al., 2008; Schalk et al., 2007; Metzger et al., 2022; Anumanchipalli et al., 2019; Ramanathan et al., 2018; Natraj et al., 2022). Given the differences in signal magnitude between low-frequency signals and *γ_H_* due to the inverse power law in neurophysiological signals, each feature data matrix was *L*_2_ normalized individually, and then concatenated together. This created a 256-dimensional data matrix over time, and this formed the online input sample to the RNN decoders.

### 6.6. Design of hDoF BCI decoders for a repertoire of stereotyped movements

#### 6.6.1. Somatotopic decoders based on spatial activity patterns

We utilized feedforward neural networks or a multi-layer perceptron (MLP) as classifiers to decode the type of movement based on purely spatial activity patterns. This network was built for each session in the initial stages of the experiment to understand the daily discernability of the movement classes. The input to the network was the 96-dimensional vector of spatially smoothed and temporally averaged grid-wise neural activity (see *Spatial feature extraction* steps in earlier sections) from individual bins during the active portions of the task. During open-loop control blocks, these bins represented ground truth labels of movement class membership. During closed-loop control blocks, irrespective of the decoder output, we assumed that subject always intended to imagine the movement associated with the cued direction (ReFit assumption (Gilja et al., 2012). Accordingly we re-labeled the decodes when using closed-loop data for decoder adaptation. To train the classifier, the data were randomly split into a training and validation set. Data augmentation techniques (detailed in subsequent sections) were also used. Using coarse grid-search, the architecture of the MLP was as follows: 3 hidden layers, 64 units per hidden layer, ReLU activation at each unit, dropout of 0.3, and weight regularization (*α* ranging from 0.2 to 0.5). Parameters of the MLP were learned via backpropagation over multiple training epochs (batch size of 32), based on cross-entropy loss with the Adam optimizer (default values (Kingma and Ba, 2014)), and with piecewise decreases in the learning rate. Early stopping was used to measure convergence during model training. Specifically, training was halted if the validation loss did not improve for six consecutive training epochs, and the saved model weights from the training epoch just before the decay in validation loss performance were used to construct the classifier. During online control, the decoder was typically run between 5Hz-10Hz. In experiments, we found that roughly equivalent performance could be achieved when using a more user-friendly training program, MATLAB’s *patternnet()* function, with its default parameters and with the same number of hidden layers, unit size, and regularization. In initial experiments, we built an MLP daily to examine our main focus on the representation of stereotyped movements. For a plug-and-play decoder, we utilized multi-day data across all three experiment types when building the MLP.

#### 6.6.2. RNN decoders based on spatiotemporal activity patterns

With the collection of longitudinal ECoG data and to improve decoding performance, we transitioned to deep learning-based decoders to better interpret spatiotemporal data (Wang et al.; Metzger et al., 2022; Moses et al., 2021; Anumanchipalli et al., 2019)). We specifically used bidirectional LSTMs (biLSTMs (Hochreiter and Schmidhuber, 1997)) a form of recurrent neural network for decoding spatiotemporal ECoG data. The architecture of the network and input data frames are detailed in Fig. S9. Briefly, rather than inputting smoothed spatial activity patterns, the input sample to the network during online control at each moment of the decoder update rate was the previous window of low-frequency and *γ_H_* features over the entire grid, resulting in input dimensions of 256 as detailed previously in *Spatiotemporal feature extraction for RNNs*. Using coarse grid-search, the parameters of the decoder were the following: the 256-dimensional sequence input was passed to two layers of stacked biLSTMs with 150 and 75 hidden units respectively with tanh and sigmoid activation functions within the LSTM cell. The hidden state of the first RNN across all the time-points formed the sequence input to the second RNN. A dropout of 30% was applied to the layers of both RNNs along with layer normalization techniques. The final hidden state of the second RNN formed the input to two linear feedforward fully connected layers. The first layer consisted of 25 units and with leaky ReLU activation. The second linear layer was the output layer of the entire network, to which the SoftMax function was applied. Data to train and validate the network was obtained from longitudinal open-loop and closed-loop ECoG data from hDoF experiments in the virtual environment. We assumed that all neural data in a given trial corresponded to the movement associated with the cued target/direction, obtaining ground truth labels for each segment. We segmented multiple temporal epochs of activity (*t* ∈ [800, 1000]) from the active portions of the task with a variable overlap between consecutive segments not exceeding 50%. In this manner, we extracted a diversity of spatiotemporal ECoG patterns for each movement, thereby improving the generalizability of the RNN. All the collected data from the virtual environment, along with data augmentation techniques (detailed in subsequent sections), were then used to train the network via backpropagation using the Adam optimizer. The batch size during training was 32, with the cross entropy loss and with piecewise decreases in the learning rate. Early stopping was used wherein training was halted if the validation loss did not improve for six consecutive training epochs. The saved model weights from the training epoch just before the decay in validation loss performance were used to construct the final RNN model. End-to-end time for signal processing, feature extraction and model training took around 2-2.5 hours on a single Nvidia GeForce RTX GPU. Having created the RNN-based decoder from the virtual environment data, we then fine-tuned the parameters of the network with neural data from operating the physical robot in the real-world environment. Specifically, we collected multiple blocks of open-loop and closed-loop center-out and stop data with the physical robot in the real-world environment. Here, the subject imagined performing the action associated with the direction of passive robot movement in 3D space. Neural features were then extracted using methods previously described and the parameters of the RNN were fine-tuned with neural data controlling the physical robot. During training, we used a much lower learning rate, a limited number of epochs, with the same validation criterion and augmentation methods detailed earlier. Using transfer learning in this manner, we were able to transition from the virtual to the real-world environment rapidly with considerably less real-world training data. During long-term PnP with the physical robot, the parameters of the RNN were kept fixed for complex reach-to-grasp and object manipulation tasks. Any fine-tuning of the fixed RNN was performed via the same procedure of collecting open-loop center-out and stop data and updating parameters with a lower learning rate. Given the complexity of hDoF control with multiple reaching and grasping phases in the real-world, we ran the RNN only at 5Hz.

#### 6.6.3. Hyperparameter tuning of hDoF decoders for PnP control

For both the MLP and the RNN, we used two hyperparameters for successful long-term hDoF PnP control across all tasks and environments. First, during real-time control, decoder decisions with probabilities less than a user-specified threshold were zeroed-out, allowing the user to disengage and not produce any output. This threshold, while typically around 0.45, varied across-sessions based on the subject’s comfort levels and experimental assessment of that day’s SNR in ECoG signals prior to starting experiments. Second, we applied a running mode filter to the running history of decodes for both the RNN and MLP i.e., the decoded action at any moment in time was the mode of the previous N decodes, with *N* ∈ (1, 5) based on task constraints, SNR and experimental assessment, and the subject’s feedback. While higher values of N introduce a slight delay between neural activity and the visual feedback of the decoded output, it also increased the single-bin level decoding accuracy. We found this to be crucial for real-time continuous hDoF control especially when executing precision trajectories with the physical Jaco arm in the real-world environment.

#### 6.6.4. Autoencoder design to understand representational similarity in somatotopic activity patterns

To provide an interpretable representation of high-dimensional spatial activity patterns for the movement repertoire, we performed dimensionality reduction using feed-forward, neural network based autoencoders. The autoencoder is a non-linear approach to estimating a latent space based on the reconstruction of the task-related spatiotemporal ECoG activity. Using coarse grid search, we designed the autoencoder to have a total of six layers, three in the encoder or compression layers and three in the decoder or reconstruction layers, including the input and output layers. The number of units in the encoder layer progressively decreased to the bottleneck layer, going from 96 (input layer corresponding to somatotopic activity patterns in all three frequency bands), to 48 (hidden layer 1), to 16 (hidden layer 2) and finally to 2 (bottleneck layer) thereby uncovering a low dimensional nonlinear manifold that captured high dimensional network-wide neural activity across the grid for the movement repertoire. The number of units in the decoder layer mirrored that of the encoder layer, progressively increasing till the full 96-dimensional input was reconstructed at the output layer. ELU activation was used throughout the network, with a dropout of 0.3. We leveraged the fact that the inputs to the network came from classes of movements with discrete labels, and thus added a parallel classifier layer on top of the bottleneck layer with as many units as the number of movements. The SoftMax activation was used for this layer. The total loss of the network was thus an additive sum of the reconstruction loss at the output layer and the cross entropy loss at the classifier layer on the bottleneck (Fig S3C). The parameters of the autoencoder were then learned via backpropagating the combined loss through the entire network using the Adam optimizer with default values. This encouraged the manifold to be interpretable while at the same time finding a low-dimensional representation that best approximated high-dimensional neural activity for the movement repertoire. To train a daily autoencoder, we used each day’s data from all open and closed-loop experiments (OL, CL1, and CL2) and from all movements during the active task states. We assumed that all neural activity during the active period corresponded to the cued movement regardless of the decoder output during real-time control. Data were randomly split into training, validation, and testing sets. Neural network parameters were learned via multiple training epochs with a batch size of 32 and with piecewise decreases in the learning rate after a set number of epochs. Early stopping was used wherein training was halted if the validation loss did not improve for six consecutive training epochs. The saved model weights from the training epoch just before the decay in validation loss performance were used to construct the autoencoder.

### 6.7. Data augmentation for hDoF BCI decoders

#### 6.7.1. For somatotopy-based decoders and autoencoders

In cases where training data were limited, augmentation was performed for better convergence of the neural network using the following strategy: the 96 dimensional training data ma-trix *X* ∈ *R^N^*^×96^, with N training samples across all 96 features, was split into its individual frequency features, resulting in individual 32-dimensional data feature matrices *X_f_* ∈ *R^N^*^×32^ for *δ, β, γ_H_*. We assumed each *X_f_* to be generated from a multidimensional Gaussian distribution and estimated its sample mean and covariance *µ_f_*, Σ *_f_*. A small non-negative number (e.g., 1 × 10^−6^) was added to the diagonal of Σ *_f_* as conditioning. We then sampled from this multidimensional Gaussian to simulate feature-specific neural data using Cholesky Decomposition i.e., 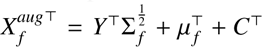 where 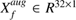 is a single vec-tor of sampled data, 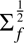 is the lower triangular Cholesky square root factor of Σ *_f_* of size *R*^32×32^, *Y* ∈ *R*^32×1^ is a column vector of unit variance, zero mean, Normal data, *µ_f_* ∈ *R*^32×1^ is the vector of feature-specific means across channels and *C* ∈ *R*^32×1^ is a column vector of zero-mean, extremely low variance Gaussian noise as conditioning for the neural network. Repeating this sampling procedure M times and then stacking all augmented feature data together subsequently resulted in an overall augmented training data matrix.

#### 6.7.2. For RNN based on spatiotemporal activity patterns

We utilized a different training data augmentation scheme, especially given the sensitivity of deep recurrent neural networks to neural data temporal diversity (Willett et al., 2021). Given trials of spatiotemporal data across all movement classes and experiment types, we augmented data by a combination of 1) adding variable DC offsets to each channel, up to ±35% of mean activity, 2) adding variable Gaussian noise along the time dimension for each channel, up to 70% of each channel’s standard deviation across time for a given trial and 3) randomly extracting variable length snippets across all *γ_H_* and low-frequency channels from random locations in trials from the original training dataset. Note that we augmented the training dataset alone and did not interfere with the validation data set. The combination of all three approaches resulted in augmenting data, that was repeatedly iterated to increase the data at a minimum by a factor of 2.5. The augmented training data was used to construct the RNN decoder using methods described in prior sections.

### 6.8. Robot control frameworks

#### 6.8.1. Input-based integrated control of continuous 3D robot end-point dynamics

We enabled continuous end-point control of the robotic arm using discrete stereotyped movements as fixed-magnitude acceleration inputs along axial directions via the following equa-tion:

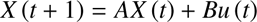

*X* (*t*) is a vector capturing the kinematic state of the robot at any time *t* and contains the following position and velocity terms in 3D space:

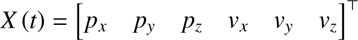

The A matrix models first-order decay dynamics where integrated velocity explains position (Gilja et al., 2012; Silversmith et al., 2021) and is given by:

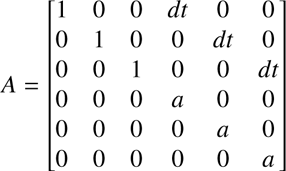

we set *a* to be 0.85 in all our experiments and *dt* is the decoding sampling interval. *u* (*t*) takes the discrete output of the BCI classifier and models it as an input or neural drive:

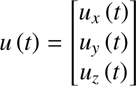

where *u_x_* (*t*) is 1,-1, or 0 depending on whether the decoded movement corresponds to the X+ve direction, X-ve direction, or neither direction, respectively. Similarly, *u_y_* (*t*) is 1, -1, or 0 and *u_z_* (*t*) is 1, -1, or 0, depending on whether the decoded movement corresponds to the +ve or -ve directions along the Y and Z axes respectively. *B* is a gain matrix to modulate the neural drive given by:

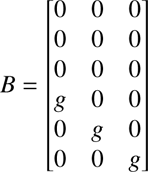

where we set *g* = 10 in all our experiments. The 7^th^ movement, ‘both middle fingers’ was used to stop the robot or perform a grasp of a target cube if the end-point was detected to be within the boundaries of the target. As this framework integrates continuous velocities based on discrete input velocity commands, we termed it IBID for input-based integrated control of continuous dynamics.

#### 6.8.2. Input-based integrated control of complex reach-to-grasp dynamics using a mode switch

To enable continuous hDoF control of both the gripper and the arm for reaching, grasping, and manipulating objects, we extended the IBID framework with a mode-switch command as detailed in Fig. 8A and 8B. The robot actions are divided into two operational modes: 1) a transport mode which allows for 3D translation of the robot gripper, and 2) a gripper mode which allows for rotation, gripper-aligned translation, and grasp open and close. Specifically, the dynamics allowed for rotation of the hand around a fixed axis in line with the gripper, and forward and backward translation of the hand to reach and withdraw from an object (Fig. 8B). The two modes utilize the same set of six imagined actions mapped to corresponding robot actions. The ‘both middle fingers’ action was used to switch between modes. To initiate the mode switch, the user must sustain the action with 80% of decodes over a 1s history, to prevent unintentional mode-switching. The space of robot actions in the gripper mode was selected to allow the user to complete the final approach and grasp within one operational mode. The user has control of three degrees of freedom: single-axis gripper rotation, gripper-aligned translation, and grasp open/close. The rotation axis is specific to the robot’s kinematic configuration and task. In the wall task, the gripper is oriented to allow for a grasp from the side of an object. Here, the user can control the “yaw” rotation around the gripper frame y-axis. The gripper-aligned translation occurs along the gripper-frame z-axis, allowing the subject to move forward and backward to grasp the object. As the subject rotates the orientation of the gripper, the translation axis rotates in world frame coordinates. In the top-down rotation task, the gripper is oriented to grasp an object from the top. Here, the user can control the “roll” rotation about the z-axis. In this case, the gripper-aligned translation along the robot-frame z-axis always corresponds to the world-frame vertical axis. As in the transport mode, the gripper mode rotation and translation are governed by first-order dynamics with the decoder output applied as an input. The dynamics are identical to the translation mode with the kinematic state now corresponding to the gripper-aligned translational and rotational velocity (rad/s). The gripper mode translation matched that of the transport mode, and for the rotation, we set *a* = 0.75 and *g* = 90 in the A and B dynamics and gain matrices respectively. The gripper state (open or close) is controlled in a discrete manner. Similar to the mode switch, the user must sustain the input corresponding to open or close with 80% of the decodes over 1s history, to prevent unintentional grasp actions.

#### 6.8.3. ReFit Kalman Filter-based continuous end-point control

We initialized and updated the parameters of a daily ReFit velocity Kalman Filter with denovo end-point continuous velocity commands using methods previously described in detail (Silversmith et al., 2021; Gilja et al., 2012; Orsborn et al., 2012) and in the supplementary section of our prior BCI study (Silver-smith et al., 2021). Here, the 3D kinematic state of the robot’s end-point is assumed to be hidden and thus has to be inferred based on continuous neural measurements, where the state is given by:

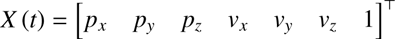

denoting 3D position and velocity terms. The last entry in the state vector accounts for the mean of neural features. It is assumed that the hidden kinematic state noisily evolves with first-order decay dynamics i.e.,

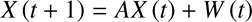

As detailed before, *A* models first-order decay dynamics with integrated position explaining velocity.

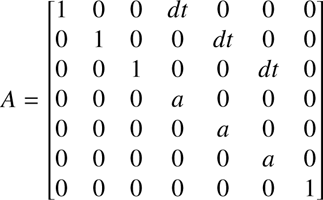

we set *a* to be 0.85 in all our experiments and *dt* is the decoding sampling interval. The noise in state evolution was given by w specific to velocity terms and modeled to be zero-mean Gaussian with known variance i.e. *W* ∼ *N* (0*, w*),

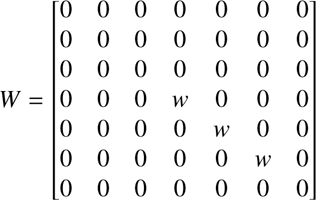

we set *w* to be 125 in all our experiments (Silversmith et al., 2021). The relationship between the state and neural measurements or observations was given by:

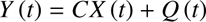

where *Y* (*t*) ∈ *R^p^*^×1^ is a vector of the *p* dimensional neural features consisting of *δ, β, γ_H_* spatial activity patterns across the grid at any time *t*. *C* ∈ *R^p^*^×7^ is the observation matrix relating neural measurements to kinematics and we modeled it to capture the state of the velocity and mean of each neural feature, with position terms set to 0. *Q* is assumed to be zero-mean Gaussian noise *Q* ∼ *N* (0*, q*) with dimensions *R^p^*^×*p*^. Given neural measurements, the kinematic state of the robot is recursively estimated as:

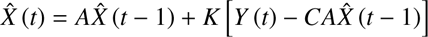

here, *K* ∈ *R*^7×*p*^ is the Kalman gain matrix that weights how well the measurements satisfy the linear model assumptions between neural and kinematic data. The estimation of the Kalman Gain is achieved via a state error covariance matrix *P* ∈ *R*^7×7^. Under the ReFit innovation, we assumed that there is only an error in estimating the velocity as the subject can perfectly infer the position of the end-effector using visual feedback. This modifies the recursive update steps in computing *P* and thereby in the solution of *K*. Full details are available in the supplementary of prior studies (Silversmith et al., 2021; Gilja et al., 2012). Given fixed parameters *A, C, Q, W* of the ReFit-KF, *K* and *P* can be recursively estimated either offline or during BCI control and typically converge rapidly to a steady-state solution (Hayes, 1996). Given that *A, W* are hand-tuned, the unknown parameters of the model are therefore only *C* and *Q*. These were initialized from open-loop trials as the subject imagined performing biomimetic hand and arm end-point commands as he passively observed the robot move to center-out targets. From the collected neural and kinematic data and hand-tuned *A, W*, the maximum-likelihood estimate of *C, Q* was computed via least-squares:

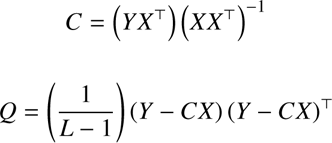

where *Y* and *X* are matrices of neural and kinematic tiled data samples across trials respectively and *L* is the total number of collected samples. Note that the velocity and mean neural terms of *C* were estimated and position coefficients were set to zero. Having estimated *C* and *Q* from open-loop data, we perform closed-loop control where the parameters were kept fixed and the subject controlled end-point position using velocity commands that were decoded by the ReFit-KF. Subsequently, we used the collected closed-loop data to update C,Q using the ReFit assumption (Silversmith et al., 2021; Gilja et al., 2012), where decoded velocity commands during online control were rotated towards the target. We adapted *C, Q* using Smooth-Batch (Orsborn et al., 2012) that exponentially weights neural data history. We chose longer half-lives (roughly 1500s on average) when updating *C, Q* as successful denovo BCI control requires slow adaptation in a 2-learner system where neural plasticity and decoder adaptation co-occur (Silversmith et al., 2021). Using smooth-batch allowed us to execute multiple fixed closed-loop control blocks (such as CL2) in a manner analogous to the discrete experiments detailed earlier.

### 6.9. Data analyses

#### 6.9.1. Imagined movement data collection for somatotopy

Trials collected during the imagined movement task (Fig. 1B) were analyzed to understand somatotopy and a representational structure in the following manner. First, all trials were z-scored to a two-minute baseline window at the start of each session wherein the subject was instructed to look at a blank screen and relax completely. The average z-scored activity during the active portion of the task (during the Go cue, Fig. 1B) over primary motor cortex channels was computed along with their confidence intervals for the movement repertoire in each of the three frequency bands. Plots of *γ_H_* activity are shown in Fig. 1C and Fig. 1D for B1 and B2 respectively. We then sought to understand channels across the grid that exhibited significant neural activity during each movement to establish cortical somatotopy. To do this, we used event-related potentials (ERPs) wherein each channel’s single-trial activity was epoched around the Go cue, from 1s before to 4s after the Go cue. Trials were then z-scored to the mean and standard deviation in the interval before the Go cue (-1000ms to 0ms) and averaged to obtain ERPs, with bootstrapped confidence intervals. A channel was deemed to be significant if any ERP response magnitude during the active task portion was outside the 95% confidence intervals in the pre-Go cue interval. This analysis was done for *δ, β, γ_H_* individually and for all movements in both B1 and B2. Finally, we used a linear support vector machine to examine the relative similarities in spatial activity patterns between all movements (Natraj et al., 2022; Fan et al., 2008). Each channel’s single-trial data during the Go cue were binned in 200ms sample intervals and then temporally smoothed over 1s to better approximate mean grid-wise spatial activity, for each of the three frequency bands. Data from the trials were then split into a training and testing set. The slack parameters in the binary linear SVMs were learned using cross-validation and by partitioning the training set into a separate validation set. Each trained binary SVM was then applied to held-out test samples, resulting in a pairwise classification accuracy indicating the similarity (accuracies closer to 0.5) or dissimilarity (accuracies closer to 1) between grid-wise spatial activity patterns. Hierarchical clustering using Ward’s criterion was performed to group movements with similar spatial activity patterns (Natraj et al., 2022; Theodoridis and Koutroumbas, 2003).

#### 6.9.2. Discrete movement decoding when mapped to cardinal 3D directions in the perspective of a virtual Jaco arm

Daily confusion matrices highlighting discriminability between movements in open-loop imagined experiments across-sessions (e.g., Fig. 2D) were computed via cross-validation. Specifically, on any given day, open-loop trials collected in the virtual robot environment were partitioned into training and test sets. A feedforward neural network (multilayer perceptron, MLP) was trained on spatial neural features from the active portions of the task in the training set. This MLP was then applied to individual bins from active task portions in the trials from the test set. Each test trial when then classified by the movement which was decoded the most (max-vote strategy) across bins. Accuracy in daily closed-loop control blocks was also determined via a max-vote strategy i.e., a given trial was classified based on which movement was decoded the most during online control; and consecutive decodes towards the cued target successfully ended the trial. Accuracies between open-loop (OL) and closed-loop control blocks (CL1, CL2) across sessions were compared using t-tests and Analyses of Variance (ANOVA) for both B1 and B2. The relative percentage improvement in classification accuracy each day during closed-loop control blocks relative to the open-loop accuracy of that day was computed for both CL1 and CL2 individually. The relative improvements in CL1 and CL2 were then statistically contrasted with each other.

#### 6.9.3. Representational structure analyses of discrete stereotyped movements

We analyzed the data from the daily discrete experiments in the virtual robot environment using autoencoders, intending to contrast the statistics of representations between the three experiment-types both within and across-sessions. Each day, an autoencoder was constructed using training and augmentation methods previously described. Once a manifold was identified, we projected high-dimensional neural data from all three experiments and all movement-classes onto the latent-space i.e., the bottleneck layer of the autoencoder. For equivalent visualization of the latent-space from two different days (Fig. 2I and 2J), a transform was applied to the second day’s latent space relative to the first. On any given day, we evaluated the pairwise separation between movements’ neural distributions in latent-space via the Mahalanobis distance given by:

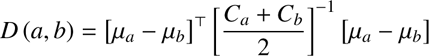

where *µ_a_* and *µ_b_* are the means of movements a and b in latent-space and *C_a_* and *C_b_* are the covariance matrices of data samples in latent-space for movements a and b respectively. We measured the pairwise Mahalanobis distances both in native high-dimensional space as well as in latent-space between movements, separately for each of the three (OL, CL1, CL2) experiments. This analysis was done for each day’s recordings for both B1 and B2 and the average pairwise distance between movements each day and for each experiment was computed. The resulting average daily distance between movements was then examined for trends across days for each of the three experiments using linear mixed effect and linear regression models. Visual inspection of data on the manifold highlighted that reductions in neural variance might be crucial in increased statistical distances between movements’ distributions during closed-loop control. As an approximate for grid-wise high dimensional neural variance, we examined the product of the eigenvalues of the covariance matrix constructed from the latent-space representation. The product of the eigenvalues denotes how much the data spread along each of its principal axes on the manifold, providing a measure of neural variance. This metric was computed for each movement and each day and the differences between OL, CL1, and CL2 were statistically contrasted using linear mixed effect models and t-tests.

To understand how our findings with low-dimensional activity patterns reflect high-dimensional data, we passed latent-space activity through the decoder layers of the autoencoder, thereby reconstructing high-dimensional neural activity. Given the design of the autoencoder and the statistics of the data, the reconstructions tended to be denoised versions of the input. We first examined the daily somatotopic similarity between the three experiments for each movement using pairwise linear correlation analyses. Here, average spatial activity was flattened into a vector for each of the three neural frequency bands and compared between experiments. Our result showed preserved somatotopy for each movement on average when transitioning from open-loop to closed-loop control. We then examined the variance of feature activity at each spatial channel of the grid. Specifically, we investigated differences in neural variance between the three experiments at each channel via its standard deviation over all trials and time-bins. Grid-wise differences in the distributions of standard deviation between the three experiments were then contrasted pairwise using Kolmogorov-Smirnov tests.

#### 6.9.4. Representational structure analyses of denovo BCI velocity commands

Similar to stereotyped movements, we built a daily autoen-coder to capture neural representations of denovo end-effector velocity commands to center-out targets in the 3D virtual workspace using a daily initialized ReFit-KF. We demarcated neural data into six labeled classes, for each of the six center-out targets in the 3D workspace. Given that continuous velocity commands span the entire workspace, we further parcellated neural data into two types to better capture target-specific neural activity. First, we built an autoencoder only for the first ∼2.5s of data across trials once online control started, captur-ing ballistic feedforward commands to various targets. Second, we extracted all bins of neural data within a trial that corresponded to accurately decoded velocities towards each target i.e., with an error less than 45 degrees; this ensures that neural data better corresponds to intended velocities towards the cued direction alone. For both types of neural data, we build a daily autoencoder across OL, CL1, and CL2 blocks using training and augmentation methods previously described. The pairwise Mahalanobis distances on the manifold between neural representations of each of the six targets were then computed. We performed statistical analyses comparing the Mahalanobis distances between both types of denovo velocity neural data and with data corresponding to discrete stereotyped representations using linear mixed effect models and t-tests.

#### 6.9.5. Representational stability analyses

Our next analyses focused on representational stability, i.e., whether neural representations were consistent across-days for the movement repertoire across all three experiment types (OL, CL1, and CL2). To simplify the comparisons in the construct of representations across-days for the movement repertoire, we relied on the autoencoder framework detailed in prior sections as a proxy for neural representations. Specifically, each day’s autoencoder was designed to best capture grid-wise spatial activity patterns across all trials and time-bins for the entire movement repertoire during both open-loop and closed-loop control. If neural representations were to be stable, then the activations at the layers of autoencoder should share some measure of similarity across-days, suggestive of an across-day stable manifold. We used the Centered Kernal Alignment method or CKA similarity metric (Kornblith et al.) to compare similarity in layer-wise activations between any two days’ autoencoders. Briefly, CKA posits that the activations at any given layer for two neural networks (when trained with similar datasets) should be invariant to either isotropic scaling or to an orthogonal transformation or both, and notably not invariant to an invertible linear transformation. In our case, if the pairwise CKA similarity across all layers between two days’ autoencoders were to be high, this would necessarily imply that the input to the network consisting of grid-wise spatial activity patterns for the repertoire share stability in their representational structure. This would imply a stable manifold. Further details on the method are available in the original paper (Kornblith et al.) detailing the various types of CKA and related metrics. For this study, we used the linear CKA similarity metric given by the following formula:

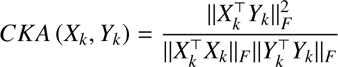

where || ∗ ||*_F_* corresponds to the Frobenius norm, *X_k_* ∈ *R^n^*^×*p*^ is the matrix of activations across the *p* units of a day *X*’s autoencoder at hidden layer *k*, *Y_k_* ∈ *R^n^*^×*p*^ is the matrix of activations across the *p* units of another day *Y*’s autoencoder at hidden layer *k*. Note that in the above equation for CKA, the activation matrices *X_k_* and *Y_k_* are mean-centered across units; we will address across-day shifts in the mean in the next section. We computed the layer-wise CKA similarity between all days’ autoencoders in both B1 and B2 in a pairwise manner. To get significance for the CKA similarities at any given layer between two autoencoders, we held the parameters of one network fixed and shuffled the weights and biases across units within individual layers of the other autoencoder before computing the CKA similarity. Similarly, the weights and biases within each layer of the first network were shuffled while holding the second network fixed, obtaining another permuted CKA similarity value across layers. The average of these two CKA similarities at all layers corresponded to one random permutation; we repeated this procedure 1000 times to get the null distribution of CKA similarities. We then assessed significance at the *α* = 0.05 level (FDR corrected for multiple comparisons (Benjamini and Hochberg, 1995)) for all pairwise and layer wise comparisons. Finally, we examined the proportion of layers between all pair-wise comparisons that were significant; this could range from 0% (no similarity) to 100% (all layers were similar). CKA analyses were done in both B1 and B2 for neural activity underlying discrete stereotyped movements and additionally in B1 for the two types of denovo velocity commands when operating the ReFit-KF. CKA values for different types of representations were contrasted via bootstrapped tests, linear statistical model and t-tests.

#### 6.9.6. Across-day drifts in centroids of neural distributions for stable well-rehearsed movements

We then assessed differences in centroids of neural activity across-days that in turn affect the location of the manifold in high-dimensional neural space. It should be noted that representational analyses so far as outlined in prior sections have been agnostic to each day’s centroid. For instance, the Mahalanobis distance between movements on the manifold is a relative distance and as such is independent of the actual location of the manifold. Similarly, the CKA metric centers activations before computing similarity patterns. We thus assessed differences between the centroids of the manifold’s day-specific neural distributions. First, data for all movements across all days were split into a training, validation, and testing set. The training and validation sets were used to build an across-day autoencoder, relying on the across-day stability of neural representations. The labels associated with individual days were used as additional class memberships when training the autoencoder and discovering the manifold using methods previously described. Held-out test data were then projected onto the latent-space, and we examined the accuracy at the classification layer on the manifold in discerning the day of recording. This method allowed us to assess shifts in centroids of neural distributions across days in latent-space for both B1 and B2. As null control, we mean-centered each day’s data to the origin and repeated the analyses.

#### 6.9.7. Discrete PnP experiments: efficacy of multi-day data and performance metrics

In general, our analyses did confirm across day shifts in the mean even with a stable construct of representations, which can be visualized as a preserved manifold whose location shifts day to day (Fig. 5E). To account for this when designing a PnP decoder (plug-and-play) we tested the efficacy of using across-day data for generalizability, with the hypothesis that the drift of stable mesoscale representations is not completely random and maybe within some tolerance of an attractor state. This tolerance may likely be within bounds that can be overcome by plasticity. To this end, we built a latent-space using a set number of days from the beginning and tested how well it captured the statistical distance between movements on held-out days. For example, we could build a manifold, using 4 days’ worth of data across all movements, and test how well it captured the representational structure for the repertoire on held-out days i.e., pairwise Mahalanobis distances between movements from days 5 onwards when projected onto the manifold. This analysis was done for cumulatively increasing days, in both B1 and B2. We examined the efficacy of using multi-day data to better capture discernability between movements using linear regression analyses. Confirmation of the efficacy of this procedure led us to use multi-day data to build a PnP MLP decoder that was tested online in two discrete experiments in the 3D virtual environment, with varying numbers of PnP days. In each experiment, there was a mode filter running on top of the decoder output, thereby accumulating decisions after the start of the active portion of the trial. The first non-zero decoded action was used to classify the accuracy of the trial based on which target/direction was cued, and the time taken to generate the decoded output was recorded. Accuracies and the time to target across blocks of trials were then used to compute bit rates, an information-theoretic measure of decoder performance (Nuyujukian et al., 2014), using the following formula:

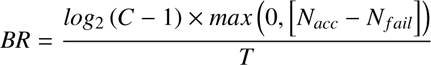

where *C* is the total number of classes or targets, *N_acc_* is the number of trials that were correctly decoded within a block of trials, *N_f ail_*is the number of trials that were inaccurately decoded within a block of trials and *T* is the total time taken within the block of trials including time taken for both accurate and inaccurate trials.

#### 6.9.8. Robot kinematics analyses

During continuous control of the robot using the IBID framework (*Input-based integrated control of continuous 3D robot end-point dynamics*) in the virtual environment, we quantified the ability of the subject to execute off-diagonal or off-axis trajectories in 3D space to distinguish from ‘etch-a-sketch’ control that is characterized by pure discrete steps in positions. We analyzed the statistics of significant off-axis control via two approaches. After collecting kinematic data towards diagonal targets placed along the 2D XY-plane the 3D workspace, we performed principal component or PCA analyses on velocity data to examine whether there was covariation; this was suggestive of continuous coarticulation as opposed to the null model wherein each PC would essentially recapitulate the X-Y axes (Fig. 7F). The statistics of coarticulation were further evaluated by examining the angular errors between the ideal diagonal velocity to the target and instantaneous decoded velocities in the IBID model across time and trials. If IBID was in agreement with the ideal velocities, then the distribution of angular errors would be non-uniform and have a mean roughly around zero degrees. However, according to the null model of etch-a-sketch control, such a distribution of angular errors will be centered about 45 degrees as discrete steps along the X and Y axes will always have a 45-degree absolute error with respect to the ideal diagonal velocity. This hypothesis was tested using the circular statistics toolbox (Berens, 2009).

#### 6.9.9. Robot reach-to-grasp metrics analyses

For the robot tasks in the real-world with the physical version of the Kinova Jaco, we assessed success rate and task completion times. In particular, success required all aspects of the task to be completed. For the wall task (Fig. 8G), this required both accurately picking up the object from the wall as well as placing it correctly at the target location on the wall and releasing the gripper. The boundaries of the object had to encompass the target location for the trial to be deemed successful. Task completion times were measured as the time when robot control first commenced to when the subject correctly placed the object and wall and released the gripper. For the reach and rotate task (Fig. 8H), we similarly measured success rates and task completion times. Here, a trial was deemed to be successful if and only if the subject completed all phases of the task correctly and rotated and placed the object at its target location after picking it up from another location. Time to task completion was measured from the time when robot control was first initiated to when the gripper released the object at the target location. For both tasks, we assessed accuracy and completion times for the duration of PnP control.

