## Supplementary Figures for "Flexible regulation of representations on a drifting manifold enables long-term stable complex neuroprosthetic control"

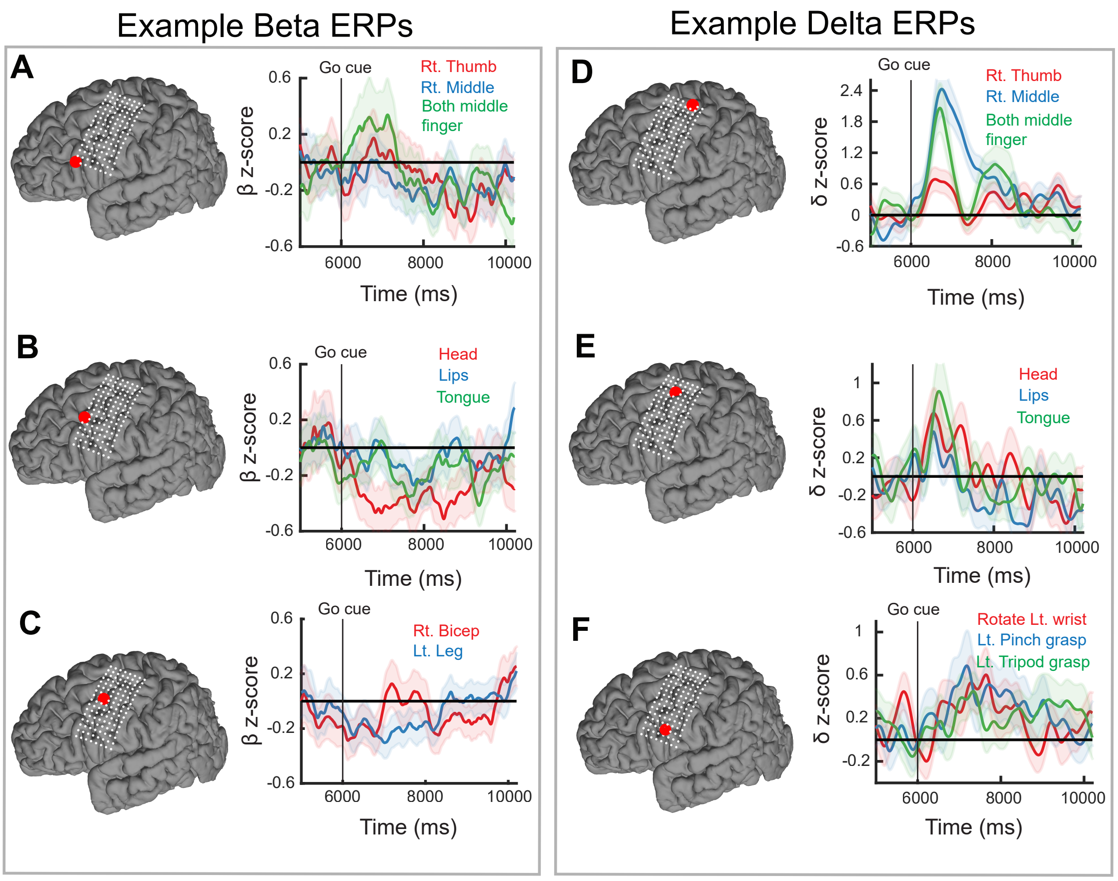


**Supplementary Figure 1:**
A-C) Example $\beta$ event related potentials over diverse electrodes (marked in red on the grid on the brain) for a diversity of imagined actions spanning the whole body. Y-axis is z-scored.
D-F) Example $\delta$ event related potentials over diverse electrodes (marked in red on the grid on the brain) for a diversity of imagined actions spanning the whole body. Y-axis is z-scored.

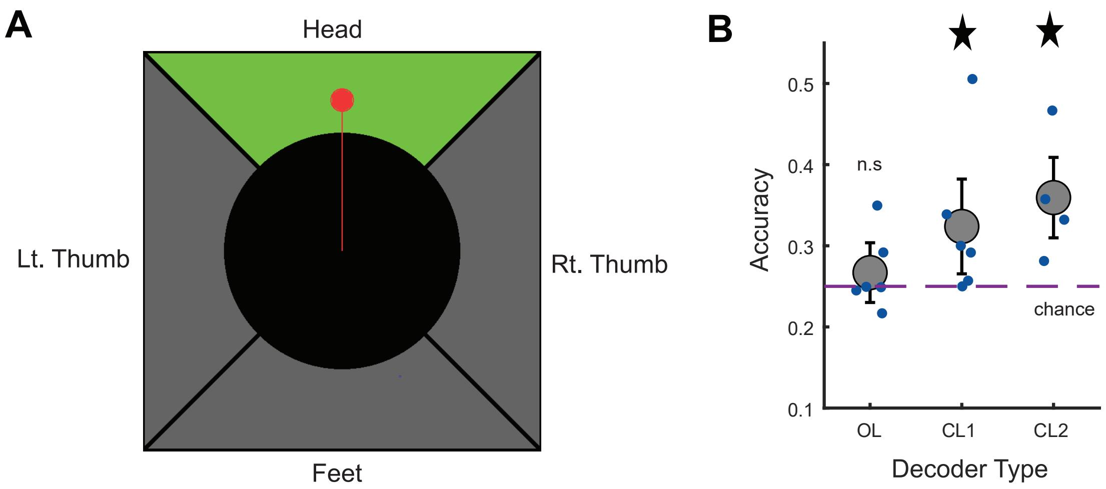


**Supplementary Figure 2:**
A) Design of the 2D experimental layout with subject B2. Each quadrant is quantized to a specific discrete well-rehearsed action and the red arrow during online trials points towards the decoded direction.
B) B2's trial-level performance in the task over multiple days during open loop (OL, cross-validated decoding accuracy) and closed-loop experimental blocks (CL1 and CL2). Each dot corresponds to a particular recording day and the mean and bootstrapped confidence intervals of average accuracy are shown. CL1 and CL2 overall had decoding accuracy significantly above chance (25% for four-class decoding, indicated with star sign).

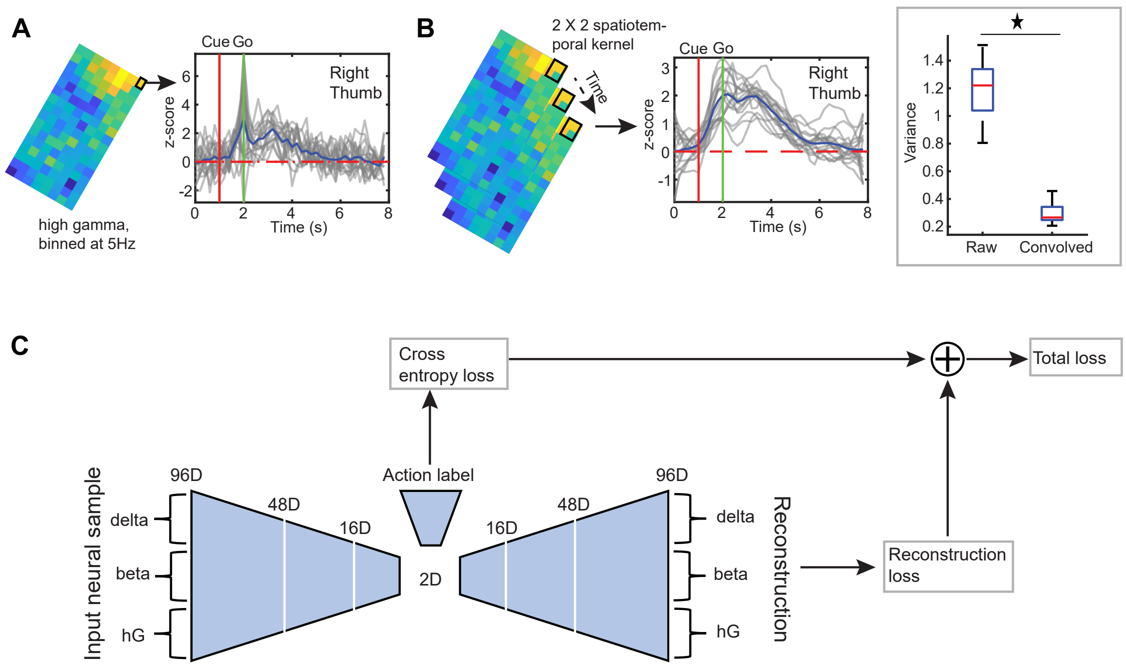


**Supplementary Figure 3:**
A) Example of single-trial binned $\gamma_H$ activity (200ms bins, grey traces, right) over a M1/S1 electrode in the hand-knob region (black square on the grid, left) during imagined right thumb movements. The mean is shown in blue and the trial markers for cue and trial start are highlighted with the vertical red and green lines respectively.
B) Example of spatiotemporal averaging of single-trial binned $\gamma_H$ activity (left, 1s temporal kernel and $2 \times 2$ spatial kernel shown on grid, and single trial grey traces, right) over the hand-knob region. Black square on the grid depicts the pseudo electrode after pooling during imagined right thumb movements. The mean activation is shown in blue and the trial markers for cue and trial start are highlighted with the vertical red and green lines respectively. Spatiotemporally smoothed (or convolved) activity during a session shows significant reductions in single trial variance across all channels over the grid as compared to raw activity and across all frequency bands ($p \leq 0.01$, star sign).
C) Design of the autoencoder for identifying a latent-space or neural manifold.

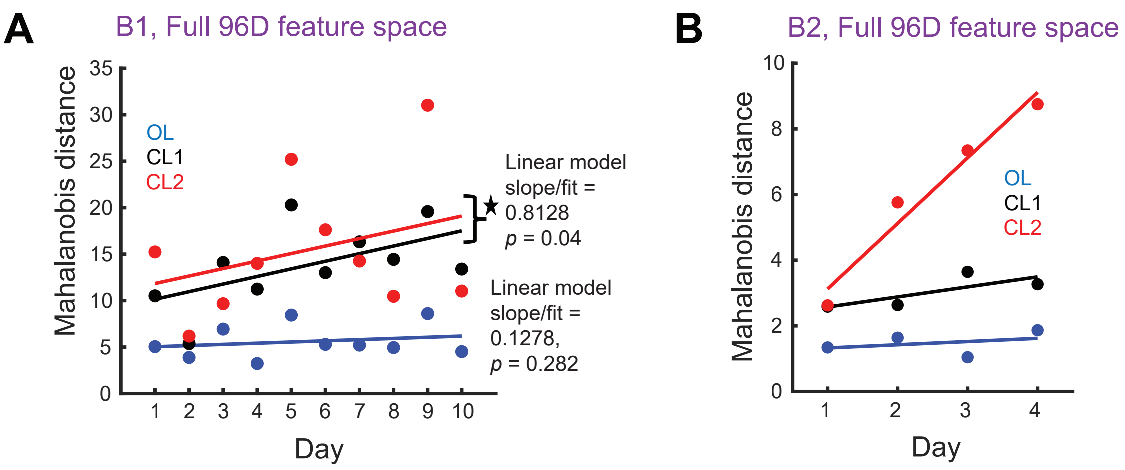


**Supplementary Figure 4:**
A) Average Mahalanobis distances across all pairwise comparisons between movements (y-axis) for each day of the experiment (x-axis) in full 96D neural feature space in subject B1. Star sign indicates significance at the $p = 0.05$ level. A mixed effect linear model revealed that CL1 and CL2 were characterized by a linear trend of increased separation between movements across days ($\beta = 0.8128$, $p = 0.0405$, *permutation-test*), in contrast to OL where the pairwise separation between movements was consistent across days ($\beta = 0.1278$, $p = 0.282$, *permutation-test*).
B) Average Mahalanobis distances across all pairwise comparisons between movements (y-axis) for each day of the experiment (x-axis) in full 96D neural feature space in subject B2.

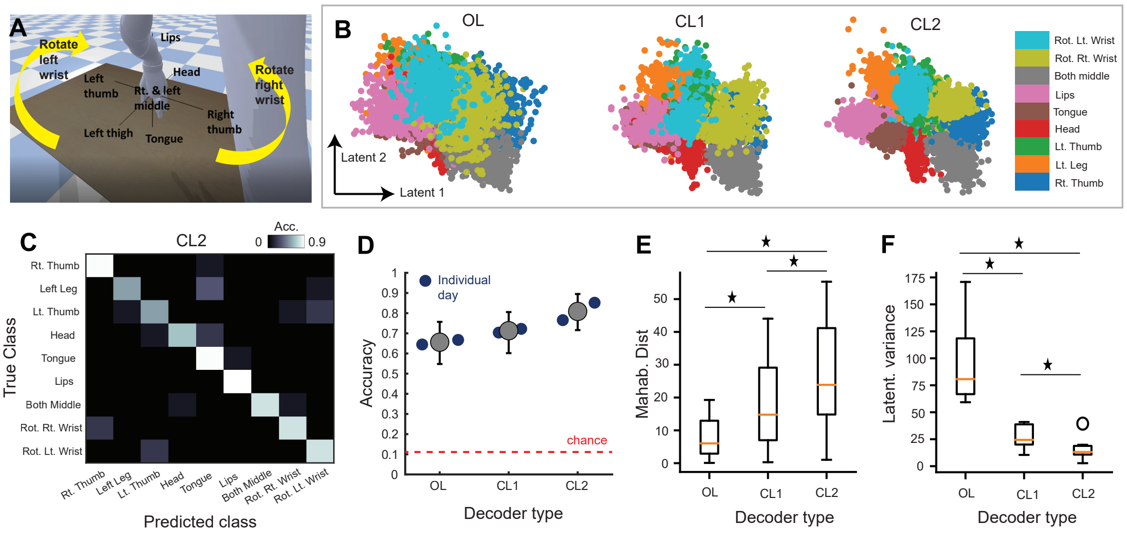


**Supplementary Figure 5:**
A) Decoding of 9 possible actions, including two new actions as commands for right and left robot arm rotation in 3D space, right wrist and left wrist rotation respectively. These two new actions in addition to the original 7 actions (main Figure 2) increase the number of discrete robot commands to 9. Similar to methods and details described in the main experiment, we collected open-loop and closed-loop BCI control data for 9 action decoding.
B) Representation of all 9 movements on the manifold or latent-space via an autoencoder. Each dot represents binned activity across all movements, sorted by the three experiment-types (OL, CL1 and CL2) and for all experimental days. The color of each data point corresponds to the particular movement.
C) Example CL2 online trial-level decoding accuracy confusion matrix for the 9 actions.
D) Mean decoding accuracies (filled circle) across all OL, CL1 and CL2 blocks, with 95% confidence intervals across trials and movements for each experimental day. Similar to the findings in Fig. 2 of the main paper, decoding accuracies tended to be higher in closed-loop control blocks (CL1, mean decoding accuracy of 71.30%, CL2 mean decoding accuracy of 80.86%) as compared to open-loop control blocks (OL, mean decoding accuracy of 65.60%).
E) Comparison of Mahalanobis distances between movements' distributions on the manifold revealed greater separation between movements on average in CL2 as compared to OL ($t\,(35) = 11.34$, $p = 2.87 \times 10^{-13}$) and CL1 ($t\,(35) = 9.066$, $p = 1.03 \times 10^{-10}$). Separation between movements was also greater during CL1 as compared to OL ($t\,(35) = 8.624$, $p = 3.5 \times 10^{-10}$). Boxplots represent all pairwise comparisons between movements for each of the three types of experiments/decoder types. Star sign represents significant difference in means between variables grouped by the horizontal line.
F) Comparison of latent variance between movements. Variance on average was consistently higher during OL as compared to CL1 ($t\,(9) = 5.522$, $p = 5.58 \times 10^{-4}$) and CL2 ($t\,(9) = 7.09$, $p = 1.03 \times 10^{-4}$), and variance in CL1 was higher than in CL2 ($t\,(9) = 2.73$, $p = 0.026$). Boxplots represent variances across all movements; outliers are depicted by open circle.

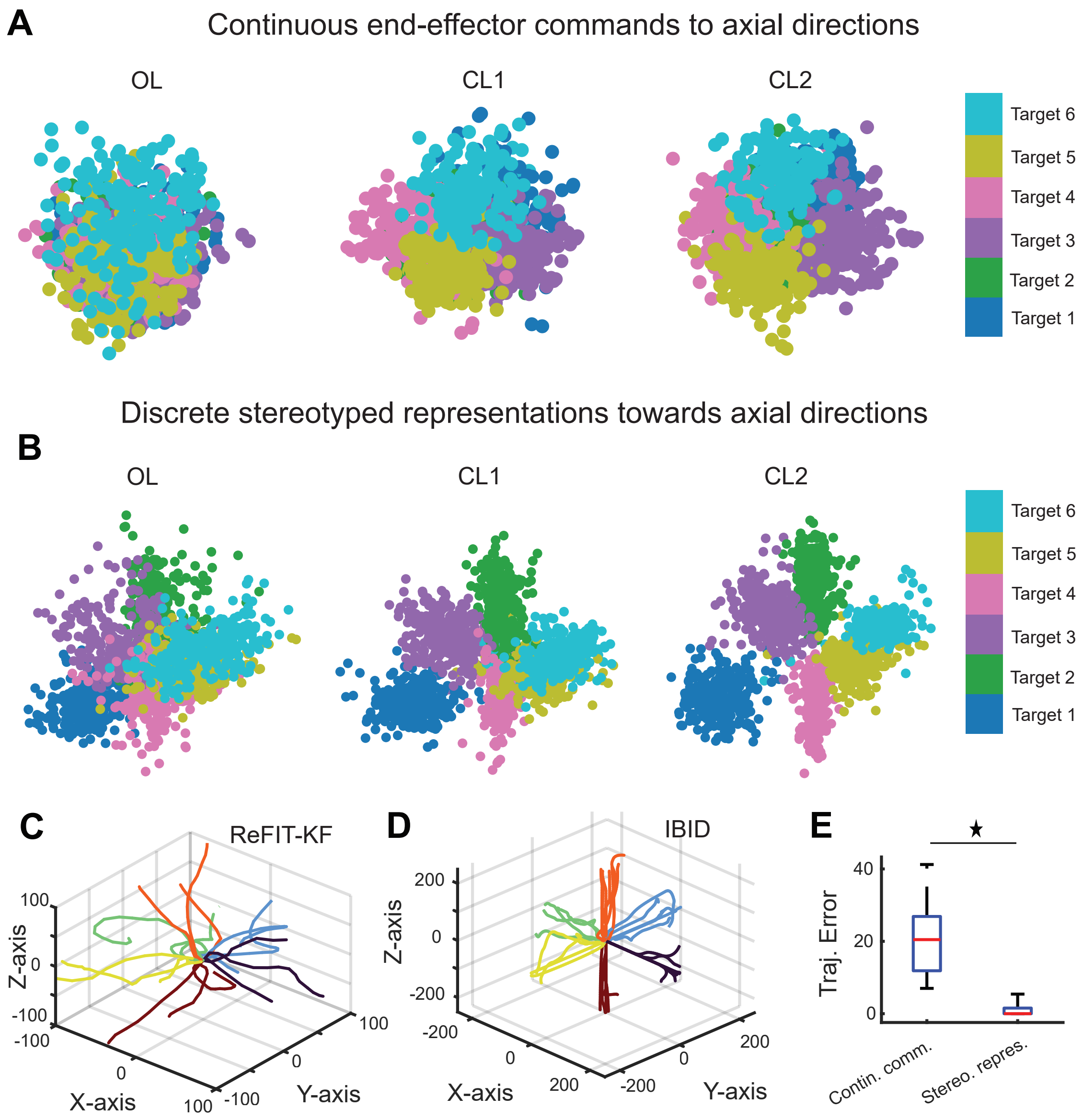


**Supplementary Figure 6:**
A) Latent-space representation of continuous 3D end-effector velocity commands (using ReFit-Kalman Filter) initialized from imagined biomimetic motor commands towards the six center out targets along axial directions of 3D space during the first 3s of the trial.
B) Latent-space representation of continuous 3D end-effector velocity commands when mapping stereotyped movement representations towards each axial direction in 3D space (see Fig. 2A) using a kinematic decoding framework detailed in Fig. 7A, for the entire trial duration.
C) Single session examples of real-time decoded trajectories during online control using the ReFit-Kalman Filter and initialized biomimetic motor commands.
D) Single session examples of real-time decoded trajectories during online control using the stereotyped representations as inputs to alter the continuous dynamics of the end-effector, i.e., Input Based Integrated control of continuous Dynamics (see Fig. 7A).
E) Boxplot showing that the ReFit-Kalman Filter decoding framework with continuous end-effector commands (initialized within-session via visual observation) produced greater errors with the ideal trajectories towards the center out axial targets as compared to stereotyped representations as discrete inputs ($t\,(63) = 12.784,\ p = 3.696 \times 10^{-19}$).

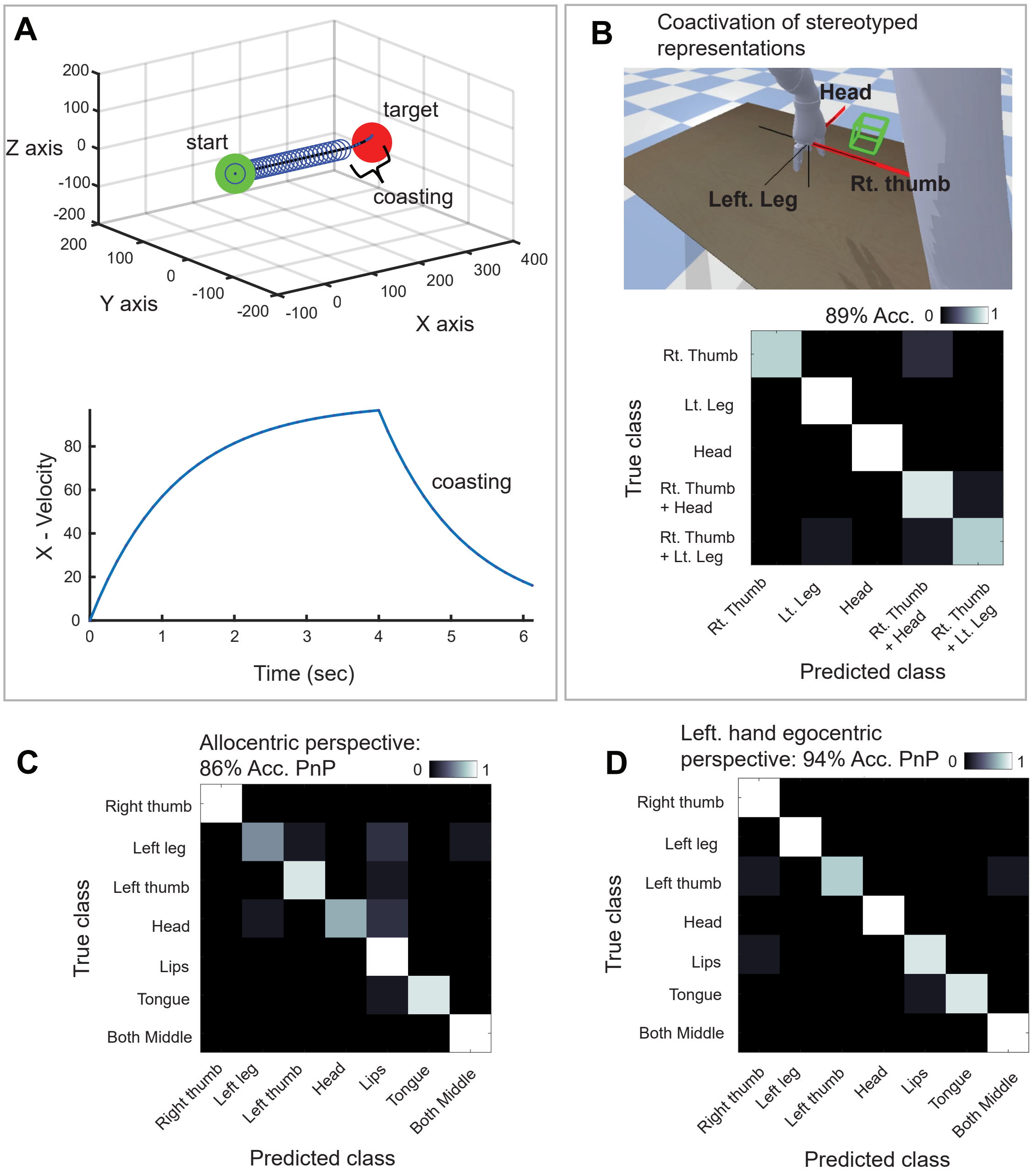


**Supplementary Figure 7:**
A) Example of 'coasting' with the input based integrated dynamics BCI controller. In the top plot, the 3D trajectory of robot position (virtual) is shown from the start to the target position. Each open circle along the trajectory depicts the decoder receiving a user input. As the robot position nears closer to the target, the user disengages and the robot 'coasts' towards the target with its own ongoing intrinsic dynamics. This is depicted in the velocity plot below, where it can be seen that the integrated velocity along the X-axis ramps up and then decays according to the first-order internal dynamics when the user disengages.
B) (Top) Design of the exemplar coactivation experiment to ascertain whether the user can successfully imagine dual movements concurrently. The green square off-axis and two axial red arrows cues the subject to coactivate two actions simultaneously. We focused on combinations with three movements relative to the right thumb. (Bottom) Trial level confusion matrix showing significant discernability between imagined single and dual coactivated movements.

**Supplementary Figure 7:**
C) Performance of the plug-and-play (PnP) decoder at the trial level during online control blocks when the targets were presented in an allocentric perspective even though the decoder was trained from a right handed ego centric perspective.
D) Performance of the plug-and-play (PnP) decoder at the trial level during online control blocks when the targets were presented in a left-handed egocentric perspective even though the decoder was trained from a right handed ego centric perspective.

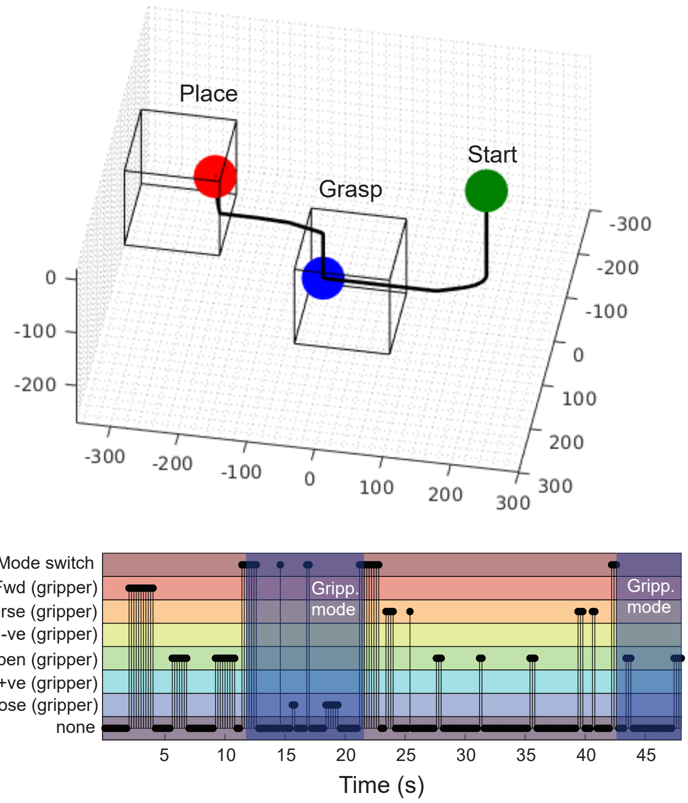


**Supplementary Figure 8:**
(Top) Robot trajectory in the virtual environment during a trial of the reach-to-grasp, transport, and place task with the input-based integrated dynamics BCI controller and 'mode-switch'. (Bottom) The temporal sequence of user inputs during the trial to drive the robot's dynamics. Periods where the user switched from transport to gripper mode and flexibly reused neural commands are shaded darker and labeled.

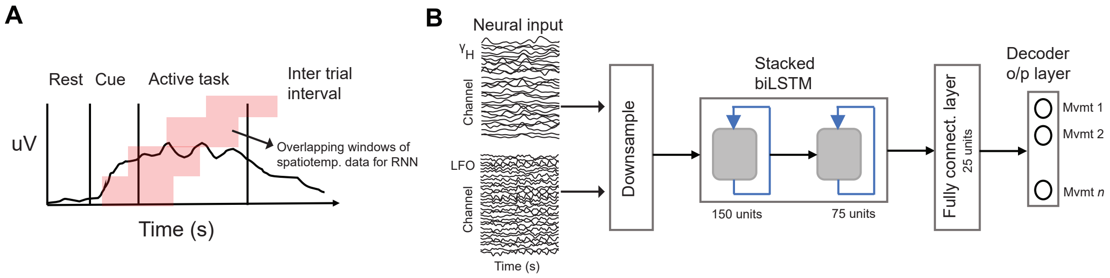


**Supplementary Figure 9:**
A) Example of how overlapping windows of raw spatiotemporal neural data were extracted for processing to train the RNN.
B) Design of the RNN; the input features, $\gamma_H$ and LFO (low-frequency, $\leq$ 25Hz) activity over all channels, were down-sampled, artifact corrected and normalized, effectively forming a 256 channel sequence input. This was fed through a stack of bidirectional LSTMs, a fully connected layer and finally a decoder layer to discern the class of movement associated with the input sample.
